# Rapid brain development and reduced neuromodulator titres correlate with host shifts in *Rhagoletis pomonella*

**DOI:** 10.1101/2021.10.13.464226

**Authors:** Hinal Kharva, Jeffrey L. Feder, Daniel A Hahn, Shannon B. Olsson

## Abstract

Host shifts are considered a key generator of insect biodiversity. For insects, adaptation to new host plants often requires changes in larval/pupal development and behavioural preference towards new hosts. Neurochemicals play key roles in both development and behaviour, and therefore provide a potential source for such synchronization. Here, we correlated life history timing, brain development, and corresponding levels of 14 neurochemicals in *Rhagoletis pomonella* (Diptera: Tephritidae), a species undergoing ecological speciation through an ongoing host shift from hawthorn to apple fruit. These races exhibit differences in pupal diapause timing as well as adult behavioural preference with respect to their hosts. This difference in behavioral preference is coupled with differences in neurophysiological response to host volatiles. We found that apple race pupae exhibited adult brain morphogenesis three weeks faster after an identical simulated winter than the hawthorn race, which correlated with significantly lower titres of several neurochemicals. In some cases, particularly biogenic amines, differences in titres were reflected in the mature adult stage, when host preference is exhibited. In summary, life history timing, neurochemical titre, and brain development can be coupled in this speciating system, providing new hypotheses for the origins of new species through host shifts.

## Introduction

Adaptation to environmental and ecological factors has been shown to play an important role in population divergence and speciation in a large number of systems (1–3). For phytophagous insects shifting to new host plants, populations potentially need to adapt their growth and development to the new host, and at the same time modulate their behavioural preference to locate that host. Furthermore, populations on novel hosts must regulate their life-history timing to coincide with the new host phenology (4–11). How these multiple events are synchronized between ancestral and novel hosts with vastly different phenologies and characteristics is an area of intense study for understanding the genesis of insect biodiversity (7,12).

*Rhagoletis pomonella* (Diptera: Tephrtidae) provides a unique opportunity to identify associations between pupal development, life history timing, and adult host choice. This species of flies originally infested fruits of the native downy hawthorn (*Crataegus mollis*) until apples (*Malus pumila*) were introduced in Eastern USA around 180 years ago (12–14). This introduction of a new host facilitated a shift in host preference from their native host, downy hawthorn, to domesticated apples, eventually leading to two host races specific to each fruit (12,13). This host shift from hawthorn to apple fruit was facilitated by two factors. First, apple and hawthorn flies are univoltine and the adults are short lived, thus each host race must emerge synchronized with the fruiting time of their host plant. Due to the earlier fruiting time of apples, apple flies initiate their overwintering dormancy, termed diapause, earlier than hawthorn flies, and emerge as adults about one month earlier as well (13). Previous work shows that the earlier seasonal adult emergence of apple host-race flies is driven solely by the timing of the termination of pupal diapause (15). The difference in adult emergence timing that drives synchronization with host fruits results in a degree of mating isolation between the apple and hawthorn races (16). Second, adults of the two host races also exhibit distinct preferences for the volatiles of their respective host fruits, which serves as an important reproductive barrier because the flies mate directly on or near the ripe host fruit (17–19). Previous studies forcing hybridisation between the host races in the laboratory showed that F1 hybrids of these two populations have altered peripheral olfactory physiology suggesting developmental abnormalities regarding their response to the host volatiles.(20). Recent studies have shown that the change in host fruit olfactory preference between the two host races does not occur at the chemoreception stage in the antenna (21–23), but rather at the first synapse of the olfactory system in the brain, the antennal lobe (24). The coupled difference in pupal diapause timing, adult host preference, and adult brain physiology in *R. pomonella* provides a unique opportunity to examine if there are corresponding differences in brain development between the races that could simultaneously impact both developmental rate and adult olfactory host choice.

Species from temperate regions like *R. pomonella* frequently use the timing of diapause to avoid the stresses of winter and also to synchronize themselves with the phenology of their hosts the next growing season (25). While diapause is often conceived of as a state of developmental arrest, it is in fact a dynamic, physiologically regulated process with defined phases of development including the diapause preparatory stage, diapause induction, diapause maintenance, and the resumption of rapid development at the end of diapause (26,27). In insects, the central nervous system (CNS) and associated endocrine glands produce neurochemicals like neurotransmitters, neurohormones, and neuropeptides that regulate diverse physiological events including the induction and termination of diapause (28). Diapause can be regulated via changes in hormone/neurotransmitter titres, receptor abundance, or regulation of specific neurochemical pathways across the stages of diapause, such as has been found with the dopamine and serotonin pathways(29–32). Apart from life history timing, many of these same neurochemicals also play important roles in insect behaviour. Biogenic amines like dopamine, octopamine, and serotonin are known to have profound impacts on adult insect behaviour across many taxa (33–36).

The dual role of neuromodulators in regulating both insect development and host-seeking behavior suggests that changes in neuromodulator titres or production at specific life stages could impact both life history timing and preference of a phytophagous insect for its host plant through changes in brain development or differentiation. Further, the aforementioned changes in neurophysiological response and preference for host volatiles between *R. pomonella* races might be rooted in changes in brain development, neuromodulation, or both. In this study, we use a variety of chemoanalytical, morphological, and immunohistochemical techniques to examine pupal-adult brain development, life history timing, and corresponding neurotransmitter levels in two closely related populations of *R. pomonella* that differ both in diapause timing and adult preference for their respective hosts. There are a number of potential neurochemical candidates and life history stages that could play a role in this host shift, and also a general lack of knowledge regarding how neuromodulators could impact this process. Therefore, our goal is to track brain development from larval to adult stages in the apple and hawthorn host races of *R. pomonella,* and identify which stages, if any, exhibit differences in neural development or neuromodulation between the host races, and how these differences are reflected in the mature adult fly, when host preference is exhibited.

## Methods

### Insect collection and maintenance

Apple and hawthorn fruits naturally infested with larvae were collected from four different sites in Michigan, USA (Grant, Fennville, Cassopolis, Lansing) in August and September 2016, and flies were reared from larvae to adulthood following previously established *Rhagoletis* husbandry methods (14). In May to August 2017, after leaving the pupae at room temperature for 15 days, they were shipped to India (with permit). This set of pupae were used to study brain development and quantification of neurotransmitters from adult flies. Eclosed adults were maintained on a diet of sugar and yeast on a 14L:10D light cycle at 25° C and 65% humidity. Post-eclosion, young flies 1-6 days old were classified as sexually immature whereas flies that were 12-14 days old were classified as sexually mature (37–39). To study the pre-winter and post-winter brain development as well as quantify neurotransmitters, a second set of pupae were collected in summer 2018. After collecting infested fruits from the above field sites, fruit were transferred to a tray with a wire mesh attached and kept in an insect-rearing room at 25±1°C, 14L:10D light cycle. Every day newly emerged pupae were collected and transferred to petri dishes with damp vermiculite, and maintained within a chamber containing a saturated potassium chloride solution to maintain ~85% relative humidity. To differentiate diapausing and non-diapausing pupae during the diapause initiation stage, four different cohorts of pupae were set aside and subjected to metabolic rate measurements once they reached either 7 or 19 days after pupariation. Other cohorts of pupae at 10 days after pupariation were transferred to a dark refrigerator at 4°C with saturated KCL solution to stimulate over-wintering diapause for six months and study post winter development until they were hand-carried to India in November 2018.

### Chemicals and Reagents

16% Paraformaldehyde EM grade (15710) was obtained from Electron Microscopy Sciences, Hatfield, PA, USA. Triton x and Bovine serum albumin were purchased from Sigma-Aldrich (Bangalore, India). All standards, ammonium acetate, Acetone, hydrochloric acid (HCl), boric acid, and reagents required for 6-aminoquinolyl-N-hydroxysuccinimidyl carbamate (AQC) synthesis, were obtained from Sigma-Aldrich (Bangalore, India). Deuterated internal standards 14 were supplied by CDN isotope (Quebec, Canada). Ascorbic acid was obtained from Himedia (Bangalore, India), and Formic acid (FA) was obtained from Fisher Scientific (Bangalore, India). Reverse-phase solid phase extraction (RP-SPE) cartridges (Strata-X, 8B-S100-TAK) were obtained from Phenomenex, Inc. (Hyderabad, India). High-purity MS grade solvents (methanol, acetonitrile, and water) were obtained from Merck Millipore (Merck Millipore India Pvt. Ltd., Bangalore).

### Pre-diapause metabolic rate measurement

Even though pupal diapause is ecologically obligate in *R. pomonella,* a small number of pupae avert diapause under laboratory conditions and directly develop into pharate adults in the prewinter period (11). To eliminate these non-diapausing individuals from our sampling before overwintering, we used a protocol adapted from Ragland *et* al 2009 & Powell *et* al 2021 to phenotype pupae as diapausing or non-diapausing by measuring g metabolic rates in the pre-winter period.

First we collected 7-day and 19-day-old pupae to measure their weight on an analytical balance with 5 μg precision (Mettler XP6, Toledo, OH, USA). Pupae were transferred to a 5 ml syringe used as a respirometry chamber for checking metabolic rate as an indicator of diapause or non-diapause status, held for 24h so adequate CO_2_ could build up in the chamber, and CO_2_ was measured on the 8th and 20th day respectively. Syringes were sealed with the plunger drawn back to produce a chamber of 1 ml internal volume. *R. pomonella* pupae were small enough to fit into one arm of the luer valve, allowing the full volume of the syringe to be injected. We also purged multiple empty syringes to serve as controls/blanks. Every 24 h, the full volume of each syringe was injected into a flow-through respirometry system consisting of a Li-Cor 7000 infrared CO_2_ analyser (Lincoln, NE, USA) with a resolution of 5 parts per million (ppm) CO_2_ interfaced to a Sable Systems International UI-2, recorded by Expedata data logging software (Las Vegas, NV, USA). The flow rate was fixed at 150 ml/min using a Sierra Instruments mass flow controller (Monterey, CA, USA). CO_2_-free air, scrubbed with a dririte-ascarite-dririte column, served as the baseline for measurements, and the system was routinely calibrated with CO_2_-free air and a certified standard mixture of 500 ppm CO_2_ in nitrogen (Airgas, Jacksonville, FL, USA). For one replicate a total of 30-40 pupae were used to calculate the metabolic rate and after that, the brain samples were dissected on cold PBS(1x) and flash frozen with liquid nitrogen. The total number of brains dissected are denoted as **n**_dissections_, and total number of brains used for staining denoted as **n**_staining_ for each sample. For further details, see Supplementary Materials, Extended Methods.

### Brain morphology and immunohistochemistry

In early November 2018, diapausing pupae were hand-carried with ice packs from Gainesville, FL, USA to Bangalore, India. After keeping pupae 6-months in the refrigerator (October 2018 - March 2019), in March 2019 all four different cohorts of pupae of the apple race were pulled out at their respective 6-month time points and left at room temperature 25°C, 14L:10D light cycle, and 65% humidity as described in Insect collection and maintenance above. Every 5 days after removal from artificial overwintering, a few pupal caps (n=5-35) were removed to observe development externally and brains were also dissected from each individual to assess brain development (Figs. 1–2). Further, pupae were individually photographed using an infinity HD camera (lumenera, model number N9033210) attached to a stereomicroscope. After that, the brain was individually dissected using 50ul of 1X phosphate buffered saline solution, PBS (PH: 7.4) on ice with fine forceps, and stored at −80°C. Four different cohorts of pupae of the Hawthorn race were pulled out in April 2019 at their respective 6 months of overwintering time points (November 2018 - April 2019) and were photographed and then dissected using the above protocol.

**Figure 1:**
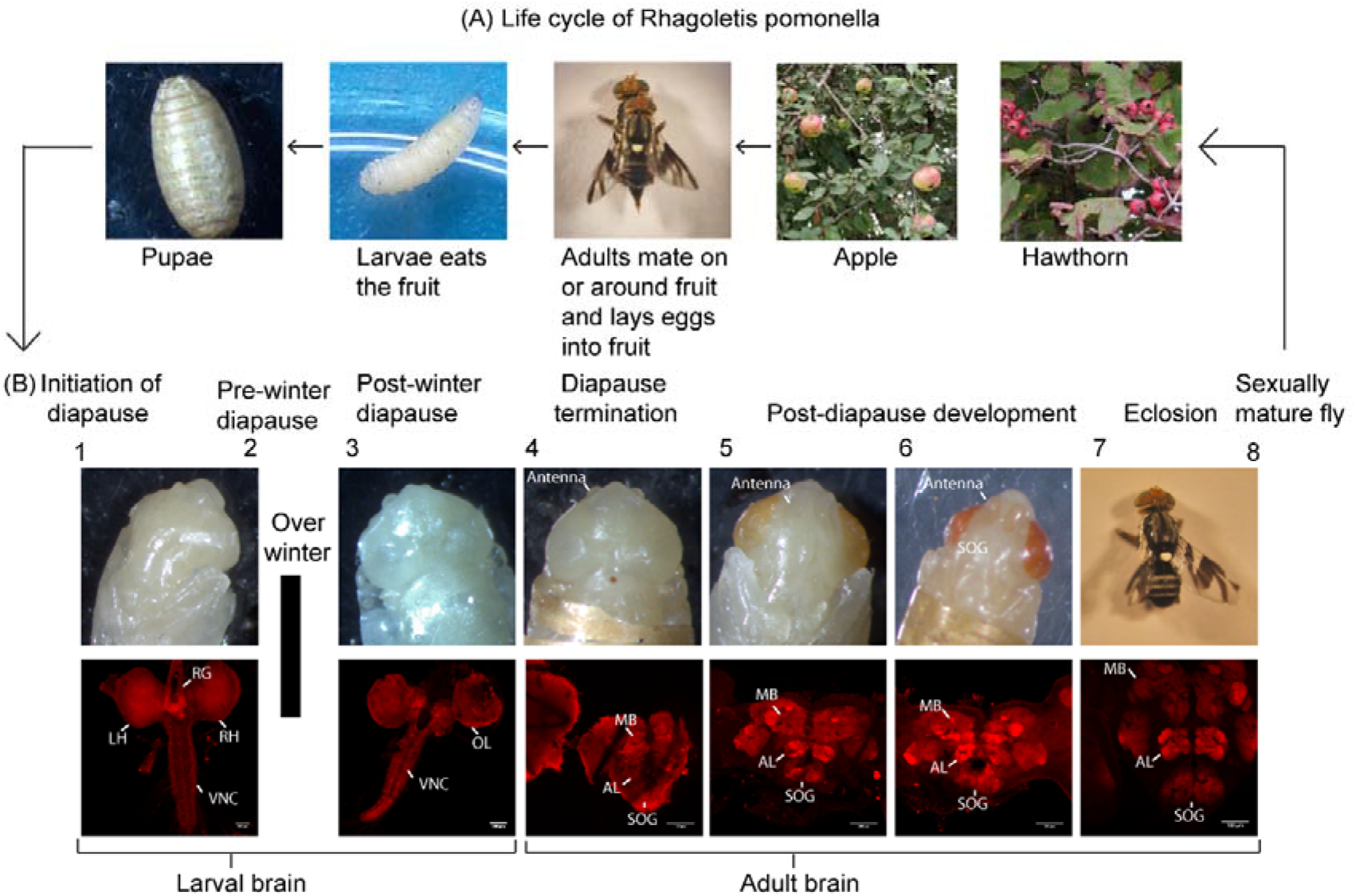
The life cycle of *Rhagoletis pomonella.* (A) Life stages of the fly from mating to oviposition from right to left. (B) Micrographic images of pupal, pharate-adult, and adult developmental stages from left to right. The upper stereomicrographs in the bottom panels show the head morphology while the lower confocal micrographs indicate the corresponding brain morphology of those stages using immunohistochemical nc82 staining across the different developmental stages (stages 1-8). The developmental stages are classified based on distinct head morphology, metabolic rate, or CNS development. Our imaging data showed that there are no morphological differences between stages 1 and 2 or between stages 7 and 8. The stages are as follows: **1**, Pre-winter diapause induction stage pupae still have high metabolic rates (Methods & Figure s1, s2 and Movie 1, apple race, stage 1); **2**, pre-winter diapause induction stage when pupae have entered metabolic depression (Figure s1, s2 Figure 1 and Movie 2, hawthorn race, stage 2); **3**, post-winter diapause maintenance phase (Figure s2 and Movie 3, apple race, stage 3); **4**, end of diapause (Figure s2. Movie 4, hawthorn race, stage 4); **5**, midway through pharate-adult development (Figure s2, Movie 5, apple race, stage 5); **6**, late pharate-adult development ((Figure s2, Movie 6, apple race, stage 6); **7**, sexually immature adult fly, less than 7 days old ((Figure s2, Movie 7, hawthorn race, stage 7); **8**, sexually mature fly, more than 12 days old; The anatomical brain regions are identified as left and right brain hemispheres (LH and RH), ring gland (RG), optic lobe (OL), ventral nerve code (VNC), the antennal lobes (AL), mushroom bodies (MB), and suboesophageal ganglion (SOG).

**Figure 2:**
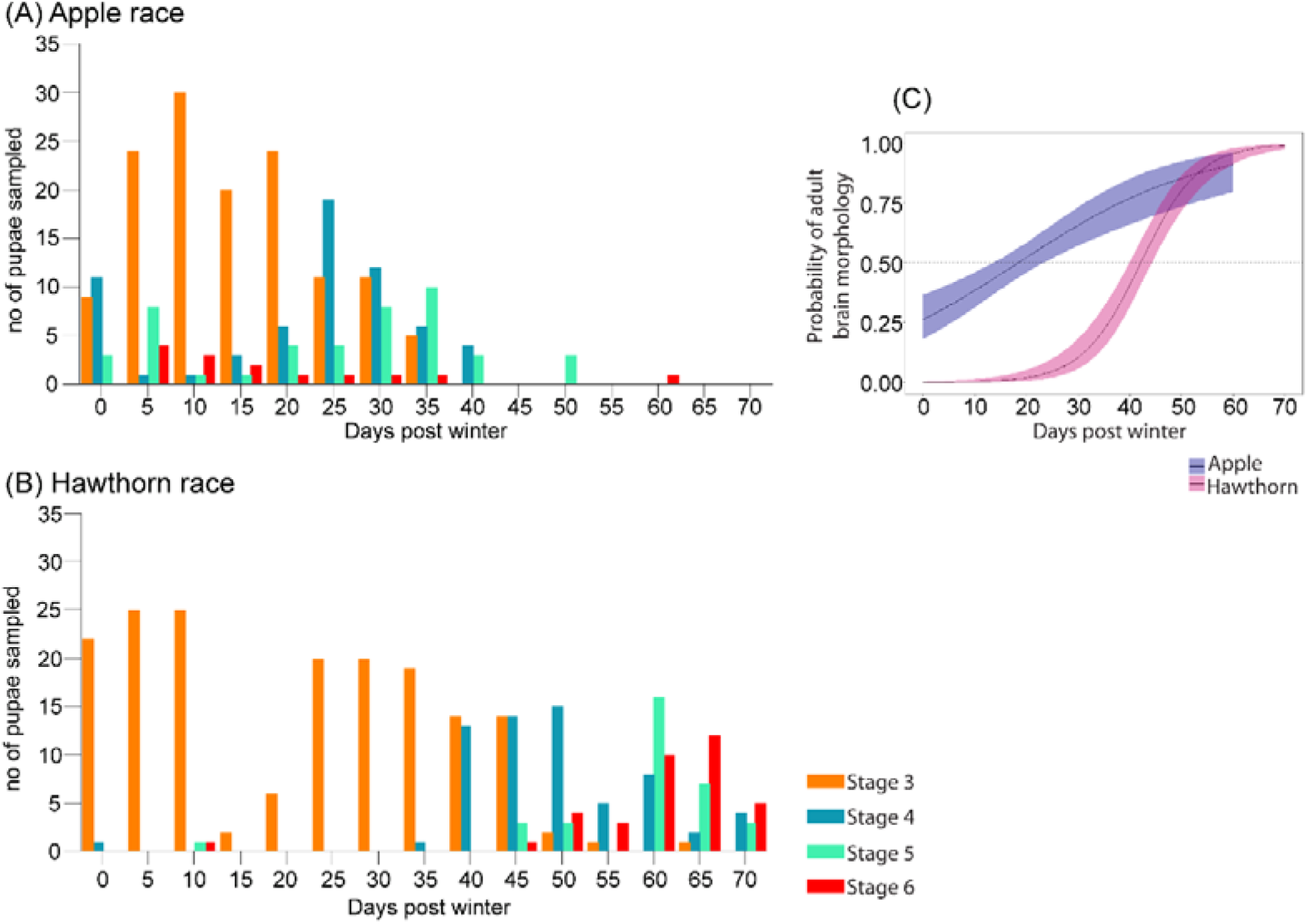
Post-winter brain development over time between the two host races. Brains were dissected from pupae and assessed every 5 days after simulated overwintering. A) Apple race, stages 3, **n**_dissections_ = 134; stage 4, **n**_dissections_ = 70; stage 5, **n**_dissections_ = 43; stage 6, **n**_dissections_ = 20; B) Hawthorn race, stages 3, **n**_dissections_ = 171; stage 4, **n**_dissections_ = 60; stage 5, **n**_dissections_ = 35; stage 6, **n**_dissections_ = 41; C) Logistic regression curves with 95% confidence intervals for the proportion of adult brain morphology vs. larval brain morphology observed in *R. pomonella* apple and hawthorn race pupae over time after artificial overwintering.

To characterize brain morphology, brain samples were dissected as mentioned above and then underwent immunohistochemical staining using a protocol adapted from those used in *Drosophila and Rhagoletis* (24,40,41). Further details can be found in Supplementary Materials, Extended Methods.

### Quantification of neurotransmitters from the R. *pomonella* brain

After morphologically determining developmental stage, brain samples were removed from −80°C storage and five brains of the same stage were pooled for each sample replicate (denoted as **n_samples_**). In one sample, only a single brain at stage 4 was observed in the hawthorn race prior to day 35 after the artificial winter. To assess the neurochemistry of this brain, we were required to pool it with other hawthorn stage 4 brains from days 35 and 40 to obtain enough material for analysis. This unique data point is shown at day 30 in Figure 4. The sample processing was performed as in Ramesh *et* al, 2019 (42–44). After the brain samples were pulled out, samples were immediately transferred to a vial containing 190 μl of Acetone (with 0.1% Formic acid, FA) and 10 μl of 1% of ascorbic acid (1.76mg/ml), followed by derivatization with 6-aminoquinolyl –N-hydroxysuccinimidyl carbamate (AQC) as in Ramesh et al 2019. (42–44). Adult fly brains were dissected individually, pooled into a group of five, and stored at −80°C with190 μl of Acetone and 10 μl of 1% of ascorbic acid. After that 10 μl of internal standard (ISTD), a mixture of all 14 neurotransmitters, (0.5ug/ml i.e. 1ng on column) was added to brain samples in screw cap vials. This mixture was sonicated for 1 min, and then homogenised using a plastic pestle. It was then immediately centrifuged (13500 rpm 4°C, 5 minutes), and the supernatant was transferred to a new tube. Simultaneously, the vials for standard solutions for calibration curves were also prepared (supplementary table s11). Serial dilutions of standard stocks were prepared with highest concentration considered on column 100% to five points including 50%,25%,12.5% and 6,25% of the maximum quantity for each of the targeted compounds, for pre-winter and post-winter brain samples. For sexually immature and sexually mature adults the standard stock was diluted to prepare 200%, 160%, 80%, 40%, 20% of the targeted compounds. After that 10 μl internal standards with 190 μl acetone (0.1% FA) and 10 μl of ascorbic acid were added to the standard solution tubes. All samples and standard tubes were dried in a speed-vac for 1hr. Apart from these, one more vial containing 16 amino acids (1 μl of 10 μg/ml) was added as an additional standard. These 16 amino acid standards helped to validate the method and retention time used for mass spectrometry (MS) every time we ran a sample. Once everything was dried in the speed-vac, 80 μl of borate buffer with 10 μl of ascorbic acid (1.76 mg/ml) was added to all tubes and vortexed. Before analysis, 10 μl of 10 mg/ml AQC (prepared in 100% Acetonitrile, ACN) was added and kept for 10 min at 55°C. After that 3ul of 100% formic acid FA was added and the tube was vortexed to stop the reaction. Tubes were subsequently kept at room temperature until all SPE columns were rinsed and cleaned with 100% methanol and 0.1% FA. After that, 500 μl of water was added to all the tubes, then vortexed and the solution was loaded on SPE columns. Columns were washed twice with 0.1% formic acid prepared with LC grade water. After that 1ml of ACN: MeOH 4:1 in 0.1% formic acid was added to the column and eluted in a new vial. All tubes were dried in a speed vac for 3hrs. Dried samples were stored at −20°C until they were run on LC-MS. Each sample was thawed and reconstituted in 50 μl of 2% ACN prepared in 0.5% FA. The LC-MS instrument method and setup are described in detail in tables s9, s10, s11, s12 figure s6 and s7. The final quantification and analyses can be found in detail in Supplementary Materials, Extended Methods.

**Figure 3:**
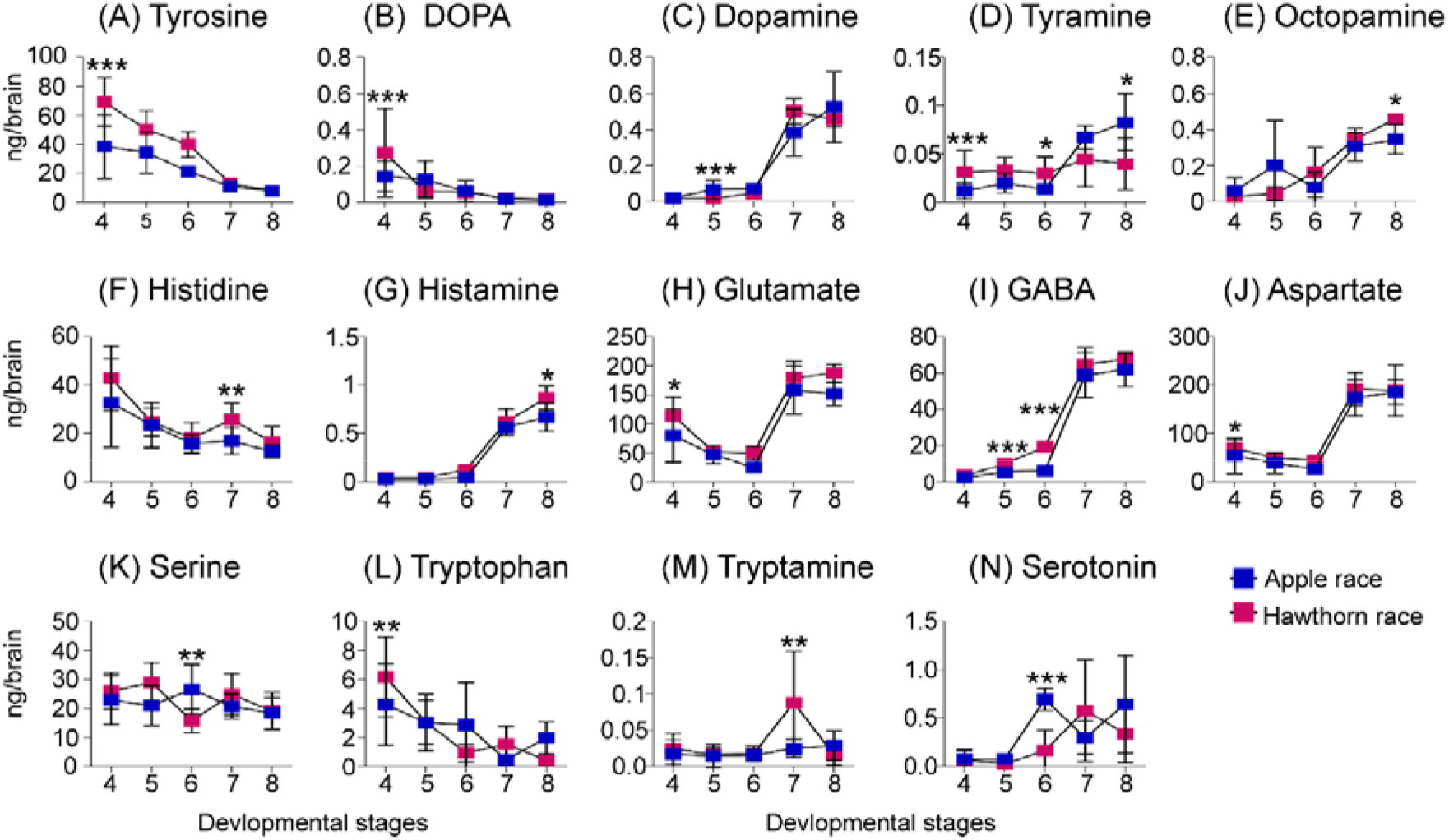
Quantification of biogenic amines and their precursors from the onset of adult brain development (stage 4) all the way up to sexually mature fly (stage 8) as defined in Figure 1 (A-N). Line graph of neurochemical titres for both host races at different developmental stages with 4-15 **n_samples_** per stage, containing a pool of five brains in each sample, symbols represent mean with SE: A) tyrosine; (B) DOPA; (C) dopamine; (D) tyramine; (E) octopamine; (F) histidine; (G) histamine; (H) glutamate; (I) GABA; (J) aspartate; (K) serine; (L) tryptophan; (M) tryptamine; and (N) serotonin; Asterisks above indicate differences between host races at the equivalent stage of brain development. P-values represented are < 0.05 *, < 0.01 **, and < 0.001 ***, linear mixed effect model, followed by Tukey’s HSD correction for multiple comparisons.

**Figure 4:**
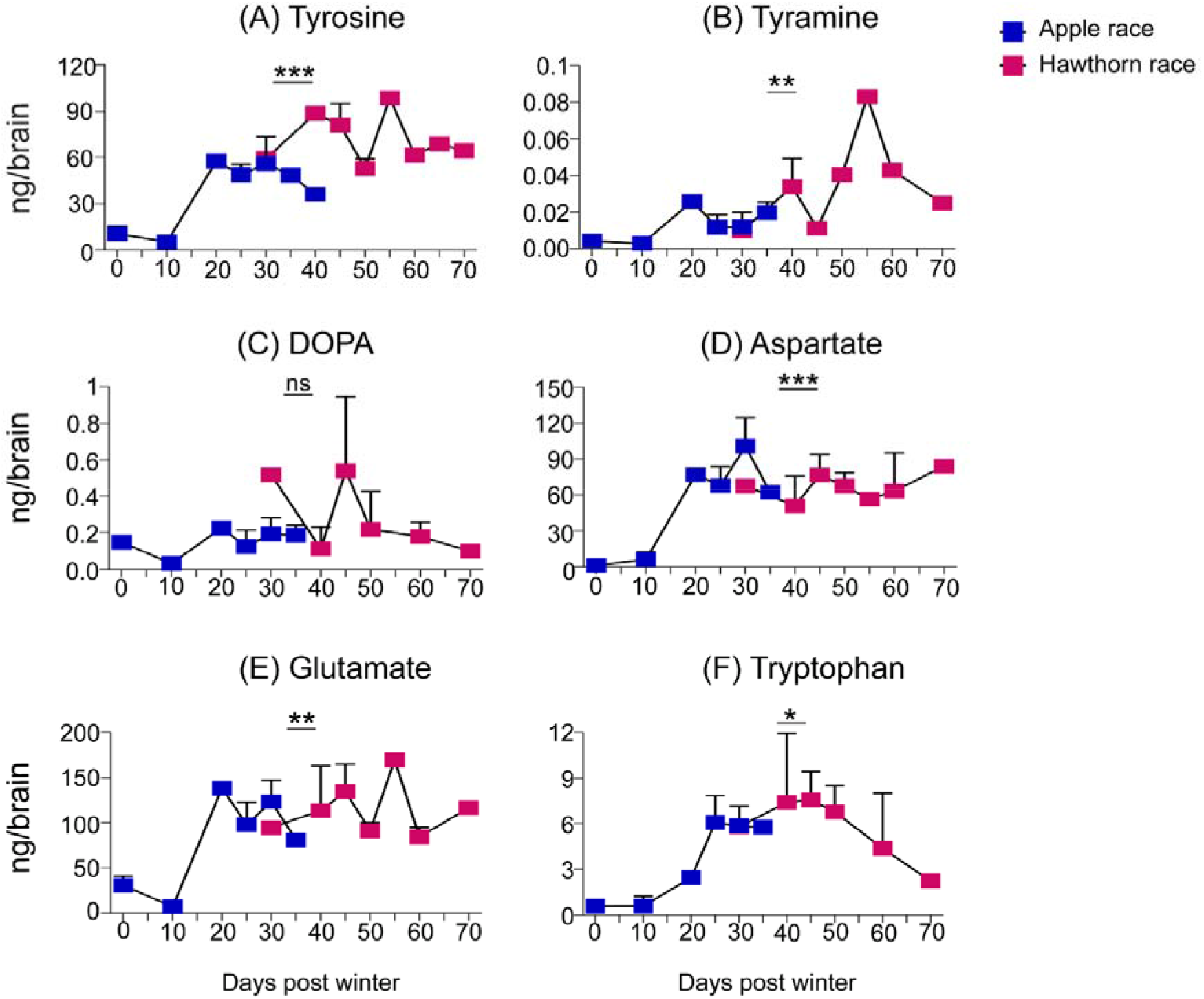
Quantification of precursor neurochemicals during onset of adult brain morphogenesis. (A-F) Line graph of neurochemical titres for both host races at different developmental stage 4 with 1-4 **n_samples_** per time point, containing a pool of five brains in each sample, symbols represent mean with SE: A) tyrosine; B) tyramine; C) DOPA; D) aspartate; E) glutamate; F) tryptophan; Asterisks above indicate significant interactions between effects of host race*days for neurotransmitter titre. P values represented are < 0.05 *, < 0.01 **, and < 0.001 ***, analysis using Two-way ANOVA, post hoc Tukey’s LSD test.

### Statistical Analysis

1) To compare the relative differences in timing of the proportion of apple vs. haw flies transitioning from pupal diapausing brain morphology to adult brain development post winter, we used a generalised linear model with a binomial, log-link function in R (V4.0.2) with ggplot, dplyr, and tidyr packed. In this logistic GLM host, days after winter was used to explain the proportion of individuals transitioning from stage 3 to stage 4 of brain development between apple and hawthorn flies, as well as to estimate 95% CIs around the logistic estimates. 2) Respiration and neuromodulator data were analysed using linear mixed effects models in R (V4.0.2) using the lmer function from the lme4 package. In the models, age and host race were used as a fixed effect whereas cohort / batch number was used as a random effect. Further, Tukey’s HSD tests with correction for multiple comparisons were performed with pairwise multiple comparisons using the lmerTest package. Satterthwaite approximations on the lmerTest package were used to test the significance of the effects. Linear Discriminant Analysis (LDA) was performed on the same transformed data using the “MASS” package in R, with “leave-one-out cross-validation” to estimate the classification accuracy of the models. 3) The interaction between host race and days on amount of precursor neurochemicals at brain development stage 4 was done using, generalised linear model, Univariate analysis of variance in SPSS V.26. In the model host race and days were used as fixed factors and quantities of each neurochemical was used as the dependent variable. Further, Tukey’s LSD tests were performed for pairwise comparisons with significance cut offs of 0.05.

## Results

### *Rhagoletis pomonella* host races exhibit distinct brain development stages during pupation

We observed external morphological and brain development in *Rhagoletis pomonella* pupae during diapause initiation, pre-winter diapause, post-winter diapause, diapause termination, pharate-adult, and eclosed adult developmental phases (Figure 1 and s2). The entire set of experiments were performed over a period of two seasons 2017 to 2019 with total of 2,091 individuals examined across both the host races. These phases were divided into 8 different stages according to changes in external head morphology, brain development, or metabolic rate (Figure 1, s1 and s2). The pre-winter diapause initiation stage 1, (8 days after puparium formation, Methods & Figure s1, s2 and Movie 1, apple **n_dissections_** = 75, **n_staining_** = 7, hawthorn **n_dissections_** = 65, **n_staining_** = 8), and the pre-winter diapause maintenance stage 2, (20 days after puparium formation, Figure s1, s2, Figure 1 and Movie 2, apple **n_dissections_** = 65, **n_staining_** = 7, hawthorn **n_dissections_** = 75, **n_staining_** = 8) were differentiated using respiration rates. Wherein pupae entering diapause exhibited higher metabolic rates 8 days after pupariation that stabilized at low levels of metabolic depression by 20 days after pupariation, when pupae were clearly in the diapause maintenance phase [Figure s1,(15,45)]. During these two pre-winter stages, brain morphology reflected that of the larval brain with a clearly identifiable ring gland and sub-oesophageal ganglion (Figure s2, A and Movie 1, 2). After a six-month artificial overwintering period at 4-5°C, we observed substantial neural differentiation and remodelling had occurred compared to pre-winter diapausing brains (stage 3, Figure s2, B and Movie 3, apple **n_dissections_** = 134, **n_staining_** = 2, hawthorn **n_dissections_** = 171, **n_staining_** = 3) in that the ring gland became disassociated and the place between the hemispheres containing the dorsal vessel became thin (c.f. stages 1 & 2; Figure s2, A, B & movie 2, 3).

Subsequent assessment of later stages also showed that adult brain (CNS) development and differentiation occurred after overwintering, at a stage during pharate-adult development where early antennal development and changes in head morphology were also observable (stage 4, Figure s2, C Movie 4, apple **n_dissections_** = 70, **n_staining_** = 4, hawthorn **n_dissections_** = 60, **n_staining_** = 1). At this stage 4, brain region boundaries were apparent, but the regions themselves remained undefined (Figure s2, C & movie 4). By stage 5, orange pigmentation had accumulated in the eyes and a transparent antenna became well developed. At this stage, the central nervous system was more defined with structures including the antennal lobe, mushroom body, and suboesophageal ganglion (Figure s2, Movie 5, apple **n_dissections_** = 43, **n_staining_** =1, hawthorn **n_dissections_** = 35, **n_staining_** = 2). The last three stages (including the sexually immature and mature eclosed adult stages) were similar in both brain and external head morphology, just with greater progression of bristle development and pigmentation of the pharate-adult cuticle (Stage 6, Movie 6, apple **n_dissections_** = 20, **n_staining_** = 1, hawthorn **n_dissections_** = 41, **n_staining_** = 1; stage 7, apple race **n_dissections_** = 45, hawthorn race **n_dissections_** = 50 Movie 7, and stage 8 apple race **n_dissections_** = 45, hawthorn race **n_dissections_** = 45, hawthorn **n_staining_** = 1 Figure s2, D).

### Apple race pupae exhibit more rapid onset of adult brain morphogenesis than the hawthorn race

To compare post-winter brain development between the two races of *R. pomonella,* both apple and hawthorn pupae were brought to room temperature (25°C) after 6 months of simulated overwintering (4-5°C) synchronized to their respective diapause initiation timing (Figure 2 A, B and Tables s1 and s2). By sampling a subset of pupae every 5 days from day 0 to day 70 after removal from artificial overwintering conditions, we observed that both host races exhibited onset of adult neurogenesis during the post-winter phases, as shown in Figure 1. In the apple race, a substantial number of pupae exhibited adult brain morphology (Stage 4-6 in Figure 1) starting from day 0 after removal from simulated winter and until day 25 when 42% of the pupae sampled exhibited adult brain morphology (57/135). Conversely, in the hawthorn race only 4/152 (3%) total pupae exhibited adult brain morphology even until day 40 (Figure 2 A, B and Tables s1, s2). Logistic regression analysis (Figure 2 C) showed that apple race individuals began initiating adult brain development significantly more rapid than hawthorn race individuals (x^2^day = 265.79, p<0.001; x^2^host = 88.04, p<0.001; x^2^day*host = 34.2, p<0.001). Diapausing apple race pupae began adult brain development approximately 24 days faster than diapausing hawthorn race pupae, with 18.34 (±0.56-0.42,95%CI) days for 50% pupae exhibiting adult brains in apple vs. 42.10 (±0.61-0.42,95%CI) days for hawthorn (Figure 2). Therefore, even though the artificial overwintering period was of the same duration for both the races, the transition from pupal to adult brain development began occurring significantly more rapid in the apple race vs. the hawthorn race. In other words, not only has the apple race shifted its overall life cycle to coincide with host phenology, but the rate of development of the pupal brain from the post winter to the initiation of adult brain morphogenesis has become more rapid.

### Neuromodulator levels in developing Hawthorn race pupal brains are generally higher than in the apple race

We next examined a total of 14 neurochemicals across 6 biochemical pathways in the developing brains of both hawthorn and apple race of *R. pomonella.* These neurochemicals included both precursor molecules and their products (Figures, 3, s3, s4 & s5). We used linear mixed models to assess both within and between host race comparisons. All statistical results have been corrected for multiple comparisons using Turkey’s HSD. In general, the titres of precursor chemicals increased first in the post-diapause stages as pharate-adult brain development began (transition from stage 3 to 4), whereas product molecule levels did not change until later stages, even as late as post-eclosion adult fly sexual maturation. Of the 14 neurochemicals examined, only four chemicals showed higher titres in the apple race, (dopamine, stage 5, p= 0.0001; serotonin, stage 6, p= 0.0001; serine, stage 6, p= 0.009). For all other molecules, the hawthorn race exhibited higher titres than the apple race. One exception is tyramine which showed higher titres in the hawthorn race at the pupal stages and higher tires at the sexually mature adult stage in the apple race (tyramine, stage 4, p= 0.0001; stage 6, p= 0.039; stage 8, p= 0.0114). These chemicals further increased in titre as *R. pomonella* development progressed, until the point of adult fly sexual maturity, when host preference first emerges. At the stage of fly sexual maturity (Stage 8, Fig. 1), product neurochemicals from two major pathways showed a difference between the host races involving histidine to histamine and tyramine to octopamine, respectively (stage 7, apple race **n_samples_** = 10, **n_dissections_** = 50; hawthorn race **n_samples_** = 10, **n_dissections_** = 50; p = 0.005; stage 8, apple race **n_samples_** = 10, **n_dissections_** = 50; hawthorn race **n_samples_** = 10, **n_dissections_** = 50; tyramine, p= 0.011; octopamine, p= 0.011; histamine, p= 0.020).

### Rapid onset of adult brain morphogenesis in the apple race corresponds to lower levels of neuromodulators

Given that in the apple race, the rate of adult brain development is significantly more rapid than the hawthorn race (Figure 2), and also exhibits lower titres of several neuromodulators, we next assessed whether the rate of development corresponded to lower levels of neuromodulators, particularly at the first stage of adult brain development (Stage 4). Figures 3, s3, s4 and s5 show that six precursor molecules including tyrosine, tyramine, DOPA, aspartate, glutamate, and tryptophan from four different biosynthetic pathways showed significant differences in titre between the host races at the first appearance of the adult brain during pupal-pharate adult metamorphic development, stage 4 in Figure 1 (stage 4, apple race **n_samples_** =14, **n_dissections_** = 70; hawthorn race **n_samples_** = 11, **n_dissections_** = 55; tyrosine, p= 0001; DOPA, p= 0.0004; tyramine, p= 0.0001; aspartate, p= 0.022; glutamate, p= 0.002; tryptophan, p= 0.002;). For each chemical, the titres were higher in the hawthorn race compared to the apple race. To compare how these precursor neurochemicals titres changed over time between the developing hawthorn and apple race pupae, we compared neurochemical titres of Stage 4 brains against the day they were sampled after winter.

A Univariate analysis of variance, UNIANOVA was conducted to examine the effects of host race and days after winter on the titres of these six precursor neurochemicals. Except for the neurochemical DOPA, there was a significant interaction between host race and day for each neurochemical titre (table s7, tyrosine, F (12,10) =10.9, P = 0.0001; tyramine, F (12, 10) = 4.95, p = 0.008; DOPA, F (12, 7) = 0.82, p = 0.632; aspartate, F (12, 11) = 10.1, p= 0.0001; glutamate, F (12, 11) = 6.47, p = 0.002; tryptophan, F (12, 10) = 4.35, p = 0.013). Therefore, neurochemical titres were significantly lower in rapidly developing apple race pupae than at later time points, when hawthorn race pupae were beginning to develop adult brain morphology. This was true for precursor chemicals even when accounting for pupae that might have been impacted by the temperature effects of development during winter itself (i.e. until day 20) (table s8, tyrosine, F (8,9) =38.14, P = 0.0001; tyramine, F (7, 10) = 4.46, p = 0.017; DOPA, F (7, 8) = 0.84, p = 0.581; aspartate, F (8, 10) = 14.25, p= 0.0001; glutamate, F (8, 9) = 25.36, p = 0.0001; tryptophan, F (7, 10) = 7.19, p = 0.003). As a result, more rapidly developing brains in apple race pupae exhibited significantly lower titres of neuromodulators (especially biogenic amine precursor molecules) than later developing hawthorn race brains, and these differences were also reflected in product molecules at the adult fly stage at which host preference is exhibited (compare stages 4 and 8 in Figures 3, s3 and s5). (37–39).

## Discussion

In this study, we have identified differences in rates of adult brain morphogenesis and levels of several neurochemicals between the apple and hawthorn host races of *Rhagoletis pomonella* across the transition from pupal diapause to post-diapause, pharate-adult development, and in the adult fly. These two closely related populations differ in both life history timing, adult host preference and are generally considered a model for incipient ecological speciation. The divergence of these two populations towards hosts with such different phenologies serves as an excellent system to examine how multiple life history events are synchronized between ancestral and novel hosts.

First, we found that while some neurogenesis occurred during the diapause maintenance phase in both host races, substantial adult brain development and differentiation was initiated only after winter and brain development progressed through several morphological stages to adult emergence. In particular, pre-diapause individuals were identical in both host race and neurochemical titres (stages 1 and 2). This indicates that any difference between the host races occurs only after diapause, and is in agreement with recent transcriptomic work showing rapid up-regulation of growth and development-related transcript genes in the apple race with shorter post-winter diapause duration as compared to the longer post-winter diapause duration of hawthorn race (46). When we dissected brains from apple and hawthorn race pupae at regular intervals after they were removed from overwintering, almost no hawthorn pupae showed brain development beyond stage 3 for at least a month after overwintering, and it took up to 50 days for the majority of hawthorn pupae to terminate diapause and transition from stage 3 to stage 4 where pharate adult neural development was apparent (Figure 2B). In contrast, some apple race pupae were already progressing to stage 4 pharate adult development as soon as they were removed from overwintering (Figure 2B). This finding indicates not only has the apple race shifted its entire life cycle to coincide with fruit phenology, but adult brain morphogenesis itself occurs roughly three weeks faster in the apple race as compared to the hawthorn race even when the overwintering period is synchronised between the host races. This change in adult brain morphogenesis is not necessarily predicted from its overall host shift, and suggests a unique phenotype in the derived apple host race.

To better understand how neurochemical signaling could be associated with the different stages of development in *R. pomonella,* we examined six major biosynthetic pathways known to impact brain development as well as behavior in insects (42,47). Our results indicate that the titres of 11 out of 14 neurochemicals were significantly reduced in the apple race pupae across multiple developmental stages as compared to the hawthorn race (Figure 3, e.g., tyrosine, tyramine, octopamine, DOPA, histidine, histamine, aspartate, glutamate, GABA, tryptophan and tryptamine). Further analysis showed that these lower titres also corresponded with earlier development of the adult brain in the apple race post diapause (Figure 4, e.g., tyrosine, tyramine, aspartate, glutamate, tryptophan).

Our main goal of this study was to link life history timing, brain development and neurochemistry in the *Rhagoletis* system by identifying particular developmental stages and neurochemicals that differ between the host races. Here, we have identified specific differences in several neuromodulators, particularly the biogenic amine pathways for octopamine and dopamine, at the first appearance of the adult brain (stage 4) and again at sexual maturity in the adult fly, when host preference is exhibited (stage 8). These differences are further coupled with a difference in developmental timing of adult brain morphogenesis between the two races. This data therefore provides specific timepoints and chemical pathways that could be assessed to determine if and how they impact life history and physiology in these populations.

For example, we show that onset of adult brain differentiation (Stage 4) also corresponds with morphogenesis and emergence of the adult antenna, and this stage is accompanied by several significant differences in neuromodulator levels between the host races. A recent study comparing olfactory neurophysiology between the apple and hawthorn races of *R*. *pomonella* identified a neuronal switch in the chemosensory system in the adult brain associated with differential host choice behaviour towards apple or hawthorn fruit (24). Such a switch in neurophysiology has similarly been observed between the Z and E strains of the European Corn Borer, *Ostrinia nubilalis,* where male preference for a particular isomer of the sex pheromone is controlled by cis-acting variation in a sex-linked transcription factor (bric à brac; bab) expressed in the developing male antenna (47). A recent study of the *Rhagoletis* brain transcriptome also shows variation in cis-regulatory elements associated with differentially expressed transcripts during diapause development and an important role of hub genes in transcriptional networks that differ during diapause development between the two host races (46). Interestingly, in *Drosophila* cis-regulatory variation in two genes involved in these pathways, tyrosine hydroxylase and dopa decarboxylase (Figure 3A), have been shown to impact neurogenesis (47). Future studies that measure the expression of enzymes involved in the production of dopamine and octopamine and selective pharmacological treatments that act as agonists and antagonists of these biogenic amines at early developmental stages in these two host races could indicate if these pathways are involved in the differentiation of these two host races.

To conclude, we have characterized the progression of neurogenesis from diapause onset to adult reproductive maturation in both the apple and hawthorn host races of *R. pomonella,* an important model for ecological speciation and diversification. We identified significantly lower neurochemical levels, particularly biogenic amines in the dopamine and octopamine pathways, in the apple race of *R. pomonella* that correspond to more rapid adult brain morphogenesis in this new host race. These differences in neurochemical titre between the races in the developing pupal stages are also apparent in the adult brains at stages when flies are reproductively mature. Because biogenic amines have been implicated to impact both pupal diapause and adult behaviour, this study offers a new hypothesis that could correlate life history timing and adult host preference through developmental differences in neuromodulation. This hypothesis must now be tested in further studies assessing enzymatic expression and pharmacological manipulation of neuromodulator levels in developing pupae. As previously suggested, connecting host preference and survival through relatively simple changes could be a widespread mechanism for generating biodiversity across phytophagous insects, contributing to the origin of the large number of species observed (24).

## Data accessibility

All data supporting this manuscript will be uploaded to Dryad upon submission.

## Authors’ contribution

HK, SBO, and DAH conceived the study and designed experiments; JLF provided the adult flies and reagents for the experiment. HK conducted experiments; HK, DAH and SBO analyzed data; HK and SBO wrote the manuscript; all authors revised and approved the manuscript.

## Acknowledgements

We would like to thank Dr, Andrew Nguyen and Dr, Chao Chen for helping with the analysis of metabolic rate. We would like to thank Dr, Cheyenne Tait for sharing *the Rhagoletis* larvae image. We also thank Khushboo Patel for helping us with the rearing of *Rhagoletis* and taking photographs of hawthorn pupal development. We thank the NCBS CIFF facility for help with confocal imaging. We thank Dr. Divya Ramesh for helping us with standardizing mass spectrometry methods and analysis.

## References

1. Schluter D. Ecology and the origin of species. Trends Ecol Evol. 2001;16(7):372–80.

2. Via S. Sympatric speciation in animals: the ugly duckling grows up. Trends Ecol Evol. 2001;16(7):381–90.

3. Berlocher SH, Feder JL. Sympatric Speciation in Phytophagous Insects: Moving Beyond Controversy ? Annu Rev Entomol. 2002;47:773–815.

4. Craig TP, Itami JK, Abrahamson WG, Horner JD. Behavioral evidence for host-race formation in Eurosta solidaginis. Evolution (N Y). 1993;47(6):1696–710.

5. Feder JL, Hunt TA, Bush L. The effects of climate, host plant phenology and host fidelity on the genetics of apple and hawthorn infesting races of Rhagoletis pomonella. Entomol Exp Appl. 1993;69(2):117–35.

6. Groman JD, Pellmyr O. Rapid evolution and specialization following host colonization in a yucca moth. J Evol Biol. 2000;13(2):223–36.

7. Itami JK, Craig TP, Horner JD. Factors Affecting Gene Flow between the Host Races of Eurosta solidaginis. Genet Struct Local Adapt Nat Insect Popul. 1998;375–407.

8. Pratt GF. Evolution of Euphilotes (Lepidoptera: Lycaenidae) by seasonal and host shifts. Biol J Linn Soc. 1994;51(4):387–416.

9. Smith DC. Heritable divergence of Rhagoletis pomonella host races by seasonal asynchrony. Nature. 1988;336(6194):66–7.

10. Wood TK, Keese MC. Host-Plant-Induced Assortative Mating in Enchenopa Treehoppers. Evolution (N Y). 1990;44(3):619.

11. Dambroski HR, Feder JL. Host plant and latitude-related diapause variation in Rhagoletis pomonella: A test for multifaceted life history adaptation on different stages of diapause development. J Evol Biol. 2007;20(6):2101–12.

12. Bush GL. Sympatric Host Race Formation and Speciation in Frugivorous Flies of the Genus Rhagoletis (Diptera, Tephritidae). Evolution (N Y). 1969;23(2):237.

13. Feder JL, Chilcote CA, Bush GL. Genetic differentiation between sympatric host races of the apple maggot fly Rhagoletis pomonella. Nature. 1988;336(6194):61–4.

14. Walsh. B. The apple-worm and the apple maggot. Am J Hortic. 1867;338–43.

15. Powell THQ, Nguyen A, Xia Q, Feder JL, Ragland GJ, Hahn DA. A rapidly evolved shift in life-history timing during ecological speciation is driven by the transition between developmental phases. J Evol Biol. 2020 Oct 1;33(10):1371–86.

16. Feder JL, Filchak KE. It’s about time: The evidence for host plant-mediated selection in the apple maggot fly, Rhagoletis pomonella, and its implications for fitness trade-offs in phytophagous insects. Entomol Exp Appl. 1999;91(1):211–25.

17. Prokopy RJ, Bennett EW, Bush GL. Mating Behavior in Rhagoletis pomonellaa (Diptera: Tephritidae) 1. Site of assembly. Can Entomol. 1971;103(10):1405–9.

18. Linn CE, Dambroski HR, Feder JL, Berlocher SH, Nojima S, Roelofs WL. Postzygotic isolating factor in sympatric speciation in Rhagoletis flies: Reduced response of hybrids to parental host-fruit odors. Proc Natl Acad Sci U S A. 2004 Dec 21;101(51):17753–8.

19. Linn C, Feder JL, Nojima S, Dambroski HR, Berlocher SH, Roelofs W. Fruit odor discrimination and sympatric host race formation in Rhagoletis. Proc Natl Acad Sci U S A. 2003 Sep 30;100(20):11490–3.

20. Olsson SB, Linn CE, Michel A, Dambroski HR, Berlocher SH, Feder JL, et al. Receptor expression and sympatric speciation: Unique olfactory receptor neuron responses in F1 hybrid Rhagoletis populations. J Exp Biol. 2006;209(19):3729–41.

21. Tait C, Batra S, Ramaswamy SS, Feder JL, Olsson SB. Sensory specificity and speciation: A potential neuronal pathway for host fruit odour discrimination in rhagoletis pomonella. Proc R Soc B Biol Sci. 2016;283(1845).

22. Olsson SB, Linn CE, Roelofs WL. The chemosensory basis for behavioral divergence involved in sympatric host shifts II: Olfactory receptor neuron sensitivity and temporal firing pattern to individual key host volatiles. J Comp Physiol A Neuroethol Sensory, Neural, Behav Physiol. 2006;192(3):289–300.

23. Olsson SB, Linn CE, Roelofs WL. The chemosensory basis for behavioral divergence involved in sympatric host shifts. I. Characterizing olfactory receptor neuron classes responding to key host volatiles. J Comp Physiol A Neuroethol Sensory, Neural, Behav Physiol. 2006;192(3):279–88.

24. Tait C, Kharva H, Schubert M, Kritsch D, Sombke A, Rybak J, et al. A reversal in sensory processing accompanies ongoing ecological divergence and speciation in Rhagoletis pomonella. Proc R Soc B. 2021;288(1947).

25. Denlinger DL, Hahn DA, Merlin C, Holzapfel CM, Bradshaw WE. Keeping time without a spine: What can the insect clock teach us about seasonal adaptation? Philos Trans R Soc B Biol Sci. 2017;372(1734).

26. Kostal V. Eco-physiological phases of insect diapause. J Insect Physiol. 2006;52(2):113–27.

27. Kostal V, Stetina T, Poupardin R, Korbelova J, Bruce AW. Conceptual framework of the eco-physiological phases of insect diapause development justified by transcriptomic profiling. Proc Natl Acad Sci U S A. 2017;114(32):8532–7.

28. Denlinger DL. Regulation of Diapause. Annu Rev Entomol. 2002;47:93–122.

29. Noguchi H, Hayakawa Y. Role of dopamine at the onset of pupal diapause in the cabbage armyworm Mamestra brassicae. FEBS Lett. 1997;413(1):157–61.

30. Andreatta G, Kyriacou CP, Flatt T, Costa R. Aminergic Signaling Controls Ovarian Dormancy in Drosophila. Sci Rep. 2018;8(1):1–14.

31. Wang Q, Hanatani I, Takeda M, Oishi K, Sakamoto K. D2-like dopamine receptors mediate regulation of pupal diapause in Chinese oak silkmoth Antheraea pernyi. Entomol Sci. 2015;18(2):193–8.

32. Zhang Q, Lu YX, Xu WH. Proteomic and metabolomic profiles of larval hemolymph associated with diapause in the cotton bollworm, Helicoverpa armigera. BMC Genomics. 2013;14(1).

33. Schulz DJ, Barron AB, Robinson GE. A role for octopamine in honey bee division of labor. Brain Behav Evol. 2002;60(6):350–9.

34. Schwaerzel M, Monastirioti M, Scholz H, Friggi-Grelin F, Birman S, Heisenberg M. Dopamine and Octopamine Differentiate between Aversive and Appetitive Olfactory Memories in Drosophila. J Neurosci. 2003;23(33):10495–502.

35. Felix A, Kromann SH, Gramsbergen J, Hansson BS, Ignell R. Dopamine regulates the mating switch in olfactory coding and preference in a female moth. 2008;1–9.

36. Kromann A. Modulation of olfactory information in the antennal lobe of Spodoptera littoralis. Modulation of olfactory information in the antennal lobe of Spodoptera littoralis. 2012.

37. Zhang A, Linn C, Wright S, Prokopy R, Reissig W, Roelofs W. Identification of a new blend of apple volatiles attractive to the apple maggot, Rhagoletis pomonella. J Chem Ecol. 1999;25(6):1221–32.

38. Nojima S, Linn C, Morris B, Zhang A, Roelofs W. Identification of host fruit volatiles from hawthorn (Crataegus spp.) attractive to hawthorn-origin Rhagoletis pomonella flies. J Chem Ecol. 2003;29(2):321–36.

39. Nojima S, Linn C, Roelofs W. Identification of host fruit volatiles from flowering dogwood (Cornus florida) attractive to dogwood-origin Rhagoletis pomonella flies. J Chem Ecol. 2003;29(10):2347–57.

40. Ibba I, Angioy AM, Hansson BS, Dekker T. Macroglomeruli for fruit odors change blend preference in Drosophila. Naturwissenschaften. 2010;97(12):1059–66.

41. Singh AP, Das RN, Rao G, Aggarwal A, Diegelmann S, Evers JF, et al. Sensory Neuron-Derived Eph Regulates Glomerular Arbors and Modulatory Function of a Central Serotonergic Neuron. PLoS Genet. 2013;9(4).

42. Ramesh D, Brockmann A. Mass Spectrometric Quantification of Arousal Associated Neurochemical Changes in Single Honey Bee Brains and Brain Regions. ACS Chem Neurosci. 2019;10(4):1950–9.

43. Rangiah K, Palakodeti D. Comprehensive analysis of neurotransmitters from regenerating planarian extract using an ultrahigh-performance liquid chromatography/mass spectrometry/selected reaction monitoring method. Rapid Commun Mass Spectrom. 2013;27(21):2439–52.

44. Natarajan N, Ramakrishnan P, Lakshmanan V, Palakodeti D, Rangiah K. A quantitative metabolomics peek into planarian regeneration. Analyst. 2015;140(10):3445–64.

45. Ragland GJ, Fuller J, Feder JL, Hahn DA. Biphasic metabolic rate trajectory of pupal diapause termination and post-diapause development in a tephritid fly. J Insect Physiol. 2009;55(4):344–50.

46. Dowle EJ, Powell THQ, Doellman MM, Meyers PJ, Calvert MB, Walden KKO, et al. Genome-wide variation and transcriptional changes in diverse developmental processes underlie the rapid evolution of seasonal adaptation. Proc Natl Acad Sci U S A. 2020;117(38):23960–9.

47. Monastirioti M. Biogenic amine systems in the fruit fly Drosophila melanogaster. Microsc Res Tech. 1999;45(2):106–21.

48. Dambroski HR, Linn C, Berlocher SH, Forbes AA, Roelofs W, Feder JL. The genetic basis for fruit odor discrimination in Rhagoletis flies and its significance for sympatric host shifts. Evolution (N Y). 2005;59(9):1953–64.

49. Unbehend M, Kozak GM, Koutroumpa F, Coates BS, Dekker T, Groot AT, et al. bric à brac controls sex pheromone choice by male European corn borer moths. Nat Commun. 2021;12(1):1–11.

