## Supplementary materials for "Rapid brain development and reduced neuromodulator titres correlate with host shifts in *Rhagoletis pomonella*"

**Supplementary Information:**

**Supplementary Data File:** This file contains extended methods, figures s1-s7, tables s1-s12 and movie 1-7

**Extended Methods:**

**Pre-diapause metabolic rate measurement**

Previous work had shown that metabolic rates dropped as pupae initiate diapause and become stabilized after they enter into the diapause maintenance phase, with a fraction of pupae not entering diapause and developing directly into pharate adults that are identifiable by their much higher metabolic rates (Ragland et al., 2009). We initially measured metabolic rates and dissected individual pupae on day 6 (n=70), day 8, day 15 (n=40), and day 20 after pupariation. This initial effort showed that day 6 individuals were still undergoing larval-pupal metamorphosis and by day 8 they showed fully developed pupal morphology. We therefore measured individuals on day 8 after pupariation to estimate metabolic rates during diapause initiation. By day 20 after pupariation, metabolic rates were lower than on day 8, showing that pupae had transitioned from diapause initiation and entered into the metabolic depression indicative of the diapause maintenance phase (Figure s1) (Powell et al., 2020; Ragland et al., 2009). Because we wished to conservatively remove any individuals that may not have been in diapause from the experiment, any individual with a metabolic rate higher than 0.025 ul CO_2_/mg/hour was eliminated from further sampling. When we dissected pupae in the low metabolic rate class indicative of being in the diapause maintenance phase at day 20 after pupariation, they clearly possessed the larval brain morphology seen in stage 2 (Fig. 1). However, when we dissected brains from pupae with metabolic rates higher than 0.025 ul CO_2_/mg/hour 20 days after pupariation these brains were in stage 4 (Fig. 1), indicating pharate-adult development had begun.

**Brain morphology and immunohistochemistry**

Briefly, dissected brains were transferred into plastic cell culture plates containing 4% paraformaldehyde (PFA), and plates were covered with aluminium foil. Brain samples were fixed in PFA for 3 hours on a rotator at 4°C under red light and subsequently washed with 0.3% Triton X prepared in 1x PBS (PTX) solution three times at an interval of 15 minutes. Brain samples were then treated with 0.1% bovine serum albumin in PTX (PBTX) as a blocking agent for 15 minutes, followed by incubation with the primary mouse anti-Bruchpilot/mAbnc82 (1:30, DSHB, University of Iowa) antibody for 48 hours on a rotator at 4°C. After two days of incubation with antibody, samples were washed four times with PTX solution at 15-minute intervals. Secondary antibody goat anti-mouse Alexa 647 (1:400, Invitrogen, Thermo Fisher Scientific, Waltham, USA) was added and brain samples were incubated for 48 hours on a rotator at 4°C. Mounting of the brain for visualization occurred on the fifth day when samples were washed with PTX 4 times, at an interval of 15 minutes at 4°C, and then mounted in 70% Glycerol on glass slides. Optical sections (512 x 512 pixels) of each brain sample with 1 µm step size, were imaged with a confocal scanning microscope (Olympus FV3000, DSS Imagetech, Bangalore) under a 10x, 20x and 40x DIC objective. Using the 3d reconstruction of *R.* pomonella brain as a guide (Tait et al., 2021) different brain regions were visualized and identified in FIJI open-source software (Schindelin et al., 2012).

**Quantification of neurotransmitters from the R. *pomonella* brain**

The level of neuromodulator per sample was calculated by a ratio of mass area of unknown sample and internal standards vs. ratio of mass area of internal standards and standards of each compound. The amount of each chemical (in ng) in the unknown sample was calculated from calibration curves. Further, outliers were identified and removed by calculating inter-quartile range (difference between Q1 and Q3 quartile) in Microsoft Excel for each age group. The lower bound was calculated by multiplying the IQR value by 1.5 then subtracting it from the Q1 data point. The upper bound was calculated by multiplying the IQR by 1.5 and adding it to the Q3 data point. In brief, the values lying 1.5 times below or above the IQR were considered outliers and removed from the further analysis. Quantities of each compound in each sample group were analysed using linear mixed model effects as mentioned above. In the model, age and host race were used as a fixed effect, whereas mass spec batch was used as a random effect. The satterthwaite approximations in the lmerTest package was used to test the significance of the effect. Further Tukey’s HSD tests were performed for pairwise multiple comparisons with significance cut offs of 0.0001,0.001, and 0.05.

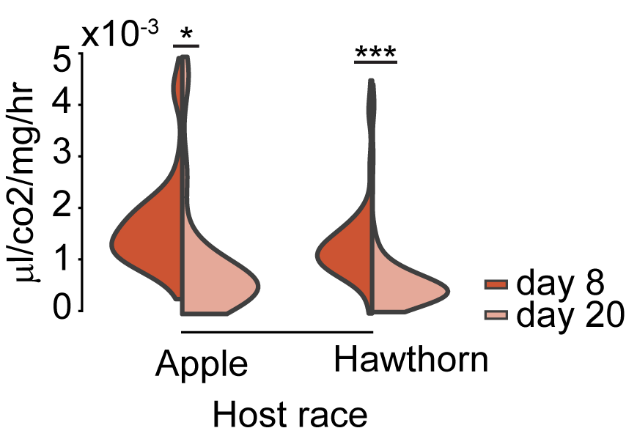

**Figure s1:** **Violin plots showing metabolic rates of pupae at two-time points for both host races**. Apple race, day 8, n=142, diapausing pupae; n=68, non-diapausing pupae; day 20, n= 145, diapausing pupae, n=62, non-diapausing pupae; hawthorn race, day 8, n=145, diapausing pupae; n=59, non-diapausing pupae; day 20, n= 152, diapausing pupae, n=17, non-diapausing pupae. Analysis performed using a linear mixed model to test for a difference in metabolic rates between day 8 and day 20. P value is significant < 0.05*, < 0.01**, < 0.001***

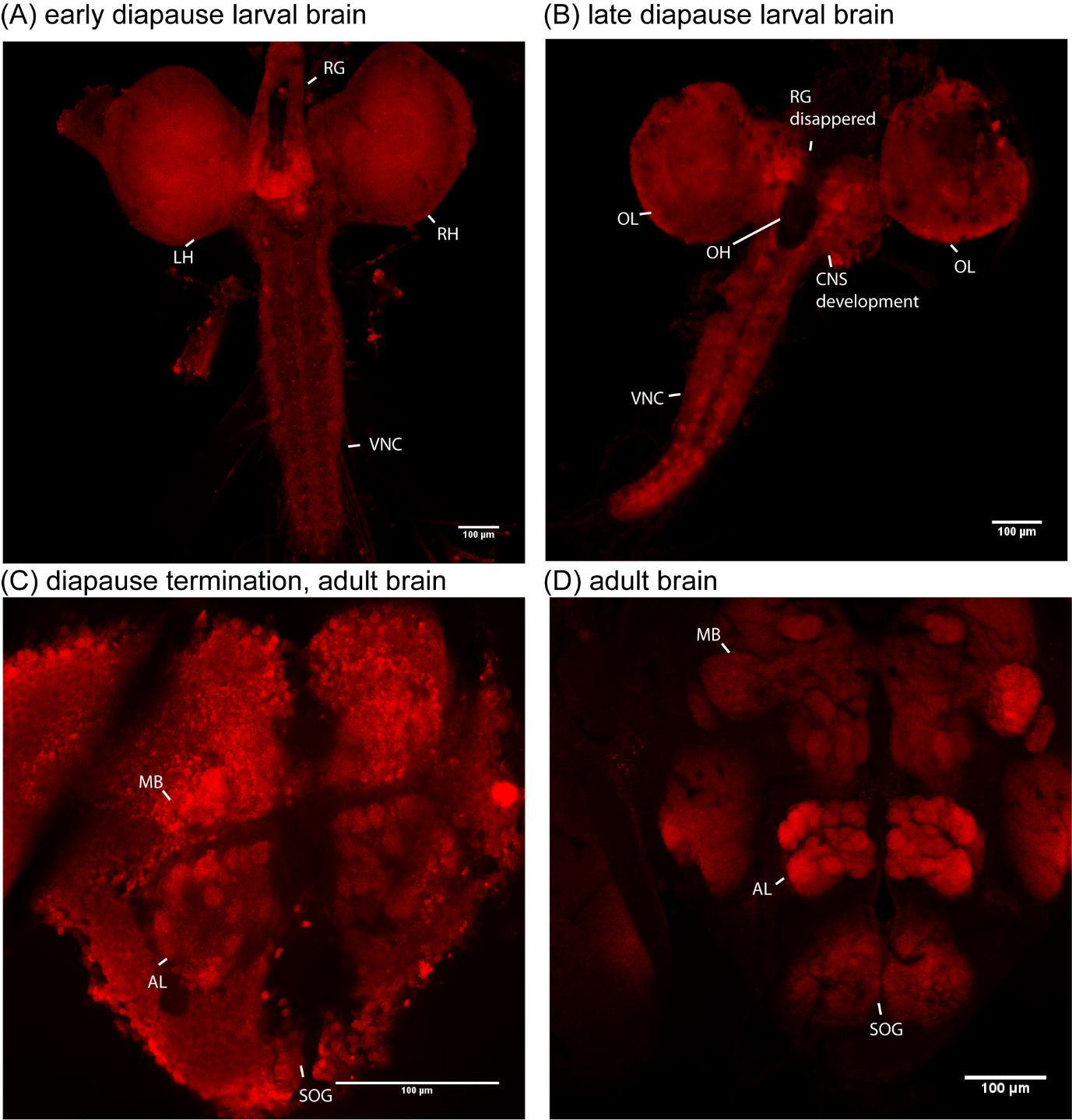

**Figure s2: Pupal brain morphology across developmental stages.** Confocal micrographs (1 µm depth) showing nc82 (red) staining of pupal brains at different stages. These are the same confocal images represented in Figure 1. The four images provide (A) an overview of the larval brain at diapause initiation (stage1, Figure 1) with two, left and right brain hemispheres (LH and RH), the ring gland (RG) and ventral nerve code (VNC), (B) the intermediate developing pupal brain at late diapause (stage 3) with clear formation of the optic lobe (OL), disappearance of the ring gland, oesophageal hole (OH) with CNS development, and VNC, (C) early adult brain (stage 4) following diapause termination with the first appearance of the boundaries of brain region like the antennal lobes (AL), mushroom bodies (MB), and suboesophageal ganglion (SOG), and (D) the adult brain (stage 7) with completely developed CNS after adult eclosion.

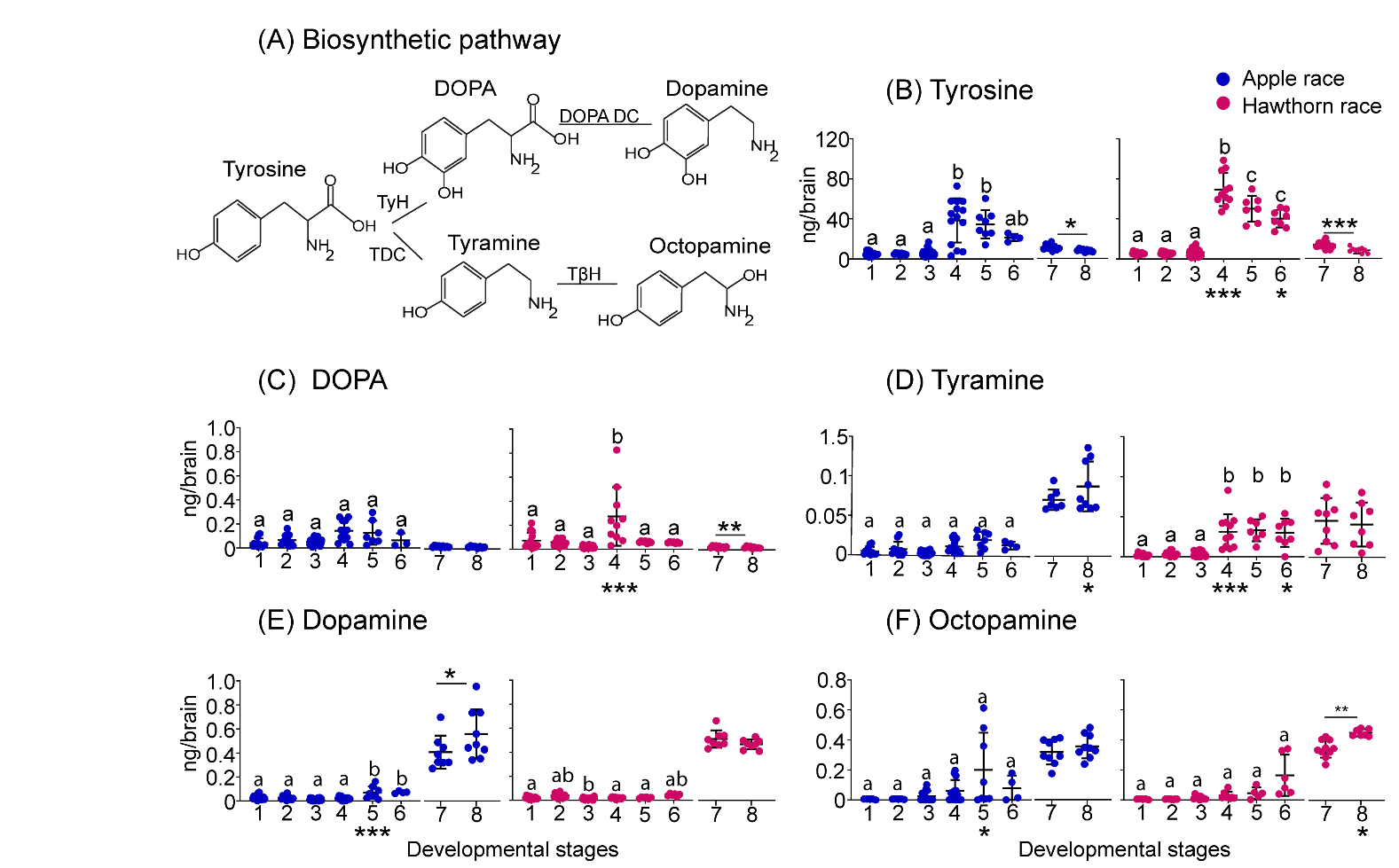
**Figure s3:** **Quantification of biogenic amines and their precursors across the different stages of development as defined in Figure 1.** (A) Biosynthetic pathways for octopamine and dopamine with enzymes, tyrosine hydroxylase (TyH), DOPA decarboxylase (DOPA DC), tyrosine decarboxylase and tyramine beta-hydroxylase (TβH); (B-F) Plots of neurochemical titres for both host races at different developmental stages with 4-15 samples per stage containing a pool of five brains in each sample, symbols represent medians with 95% CI: B) tyrosine; (C) DOPA; (D) tyramine; (E) dopamine; (F) octopamine. Letters below x-axis highlight specific developmental stages: N, onset of adult brain neurogenesis, D, development and differentiation of adult brain associated with entering the pharate-adult phase, and E, adult eclosion. Letters above each stage indicate similar titres of mentioned neurochemicals between pupal and pharate-adult stages within each host race (P > 0,05), and asterisks indicate differences in neurochemical titre between the two adult fly stages – pre-maturation and after sexual maturation. Asterisks below the x-axis indicate differences between host races at the equivalent stage of brain development. P-values represented are < 0.05 *, < 0.01 **, and < 0.001 ***, linear mixed effect model, followed by Tukey’s HSD correction for multiple comparisons.

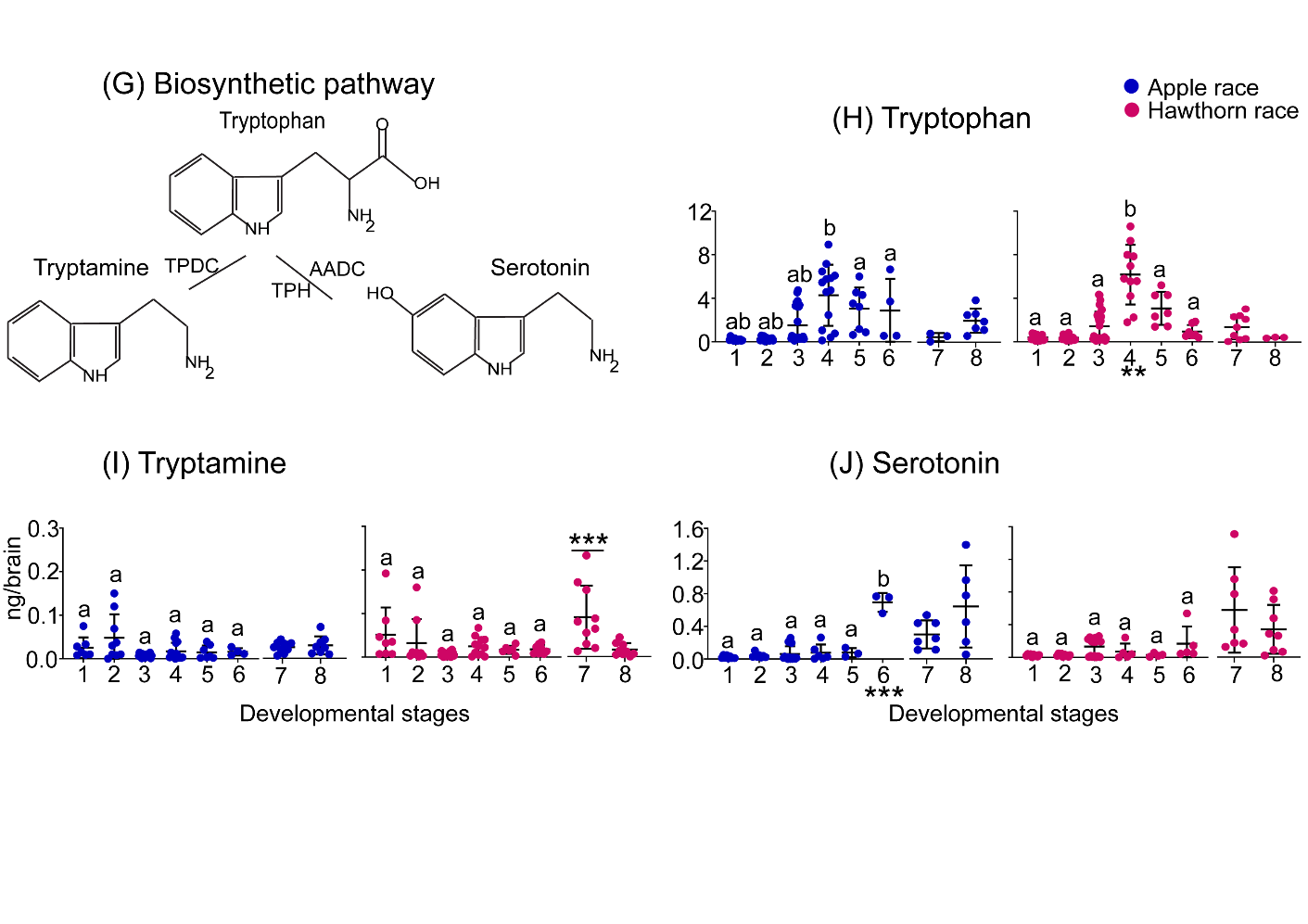
**Figure s4**: **Quantification of biogenic amines and their precursors across the different stages of development as defined in Figure 1.** (G) Biosynthetic pathway for serotonin with enzymes tryptophan decarboxylase (TPDC), tryptophan hydroxylase (TPH) and amino acid decarboxylase (AADC); (H-J) Scatter plots of neurochemical titres for both host races at different developmental stages with 4-15 samples per stage containing five brains each, symbols represent median with 95% CI: (H) tryptophan; (I) tryptamine; (J) serotonin. Lettering, asterisks, and statistical comparisons as in Figure s3.

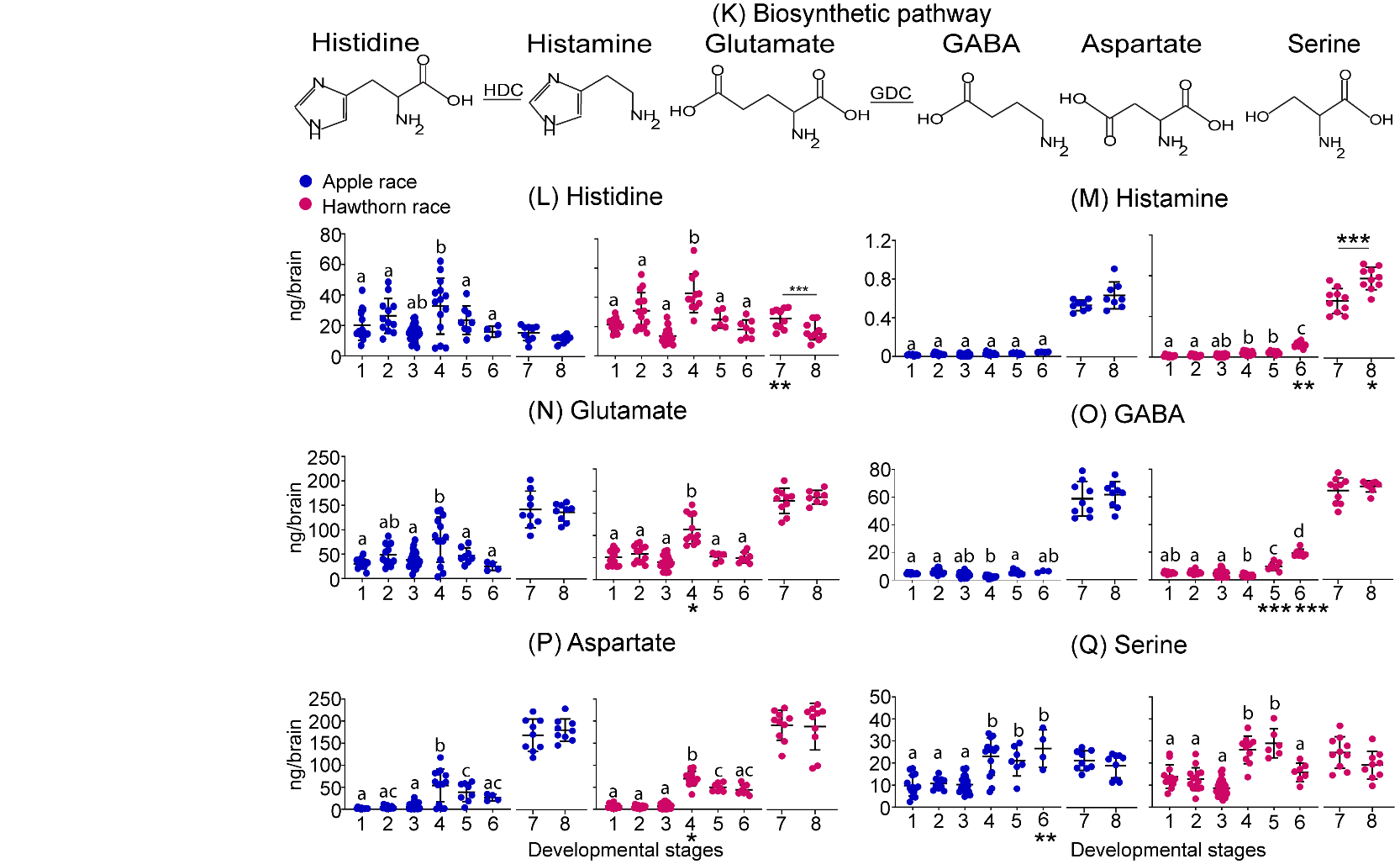

**Figure s5:** **Quantification of biogenic amines and their precursors across the different stages of development as defined in Figure 1.** (K) Biosynthetic pathway for histamine, GABA, aspartate and glutamate with enzymes histidine decarboxylase (HDC) and glutamic acid decarboxylase (GDC); (L-Q) Scatter plots of neurochemical titres for both host races at different developmental stages with 4-15 samples per stage containing five brains each, symbols represent median with 95% CI: (L) histidine; (M) histamine; (N) glutamate; (O) GABA; (P) aspartate; (Q) serine. Lettering, asterisks, and statistical comparisons as in Figure 3.

**A)**

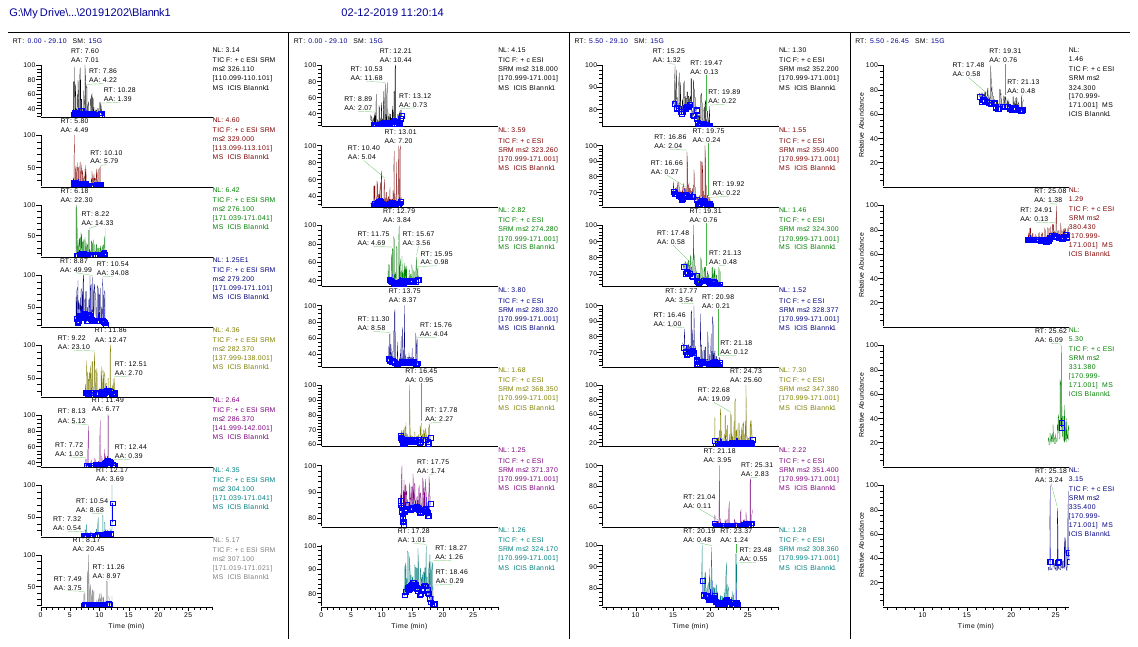

**B)**

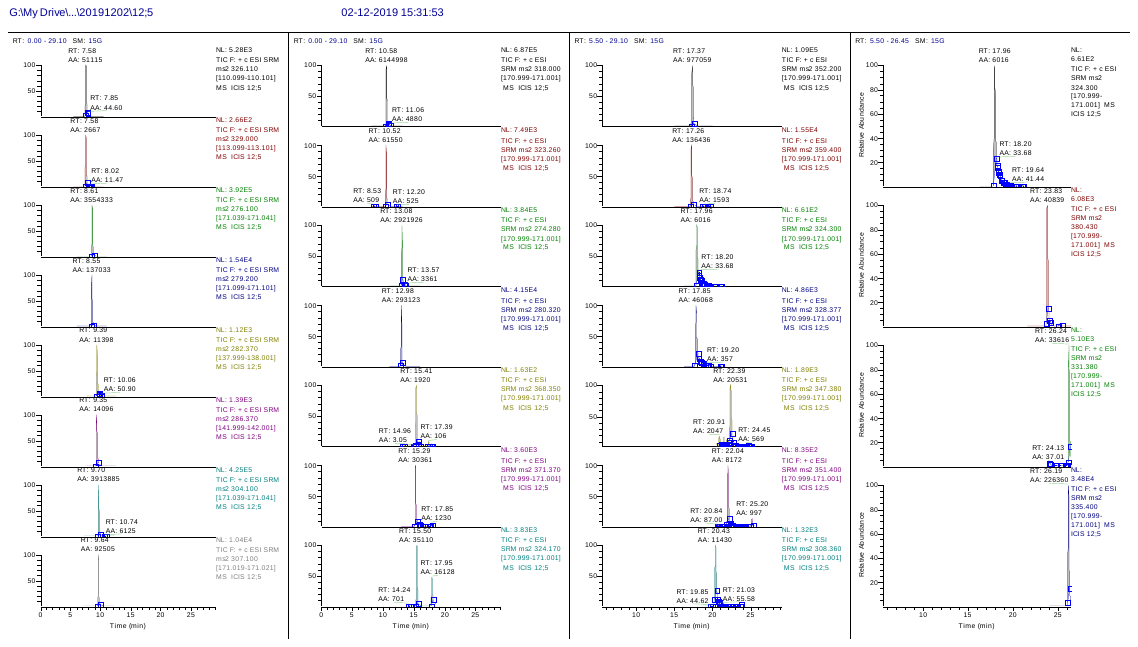

**C)**

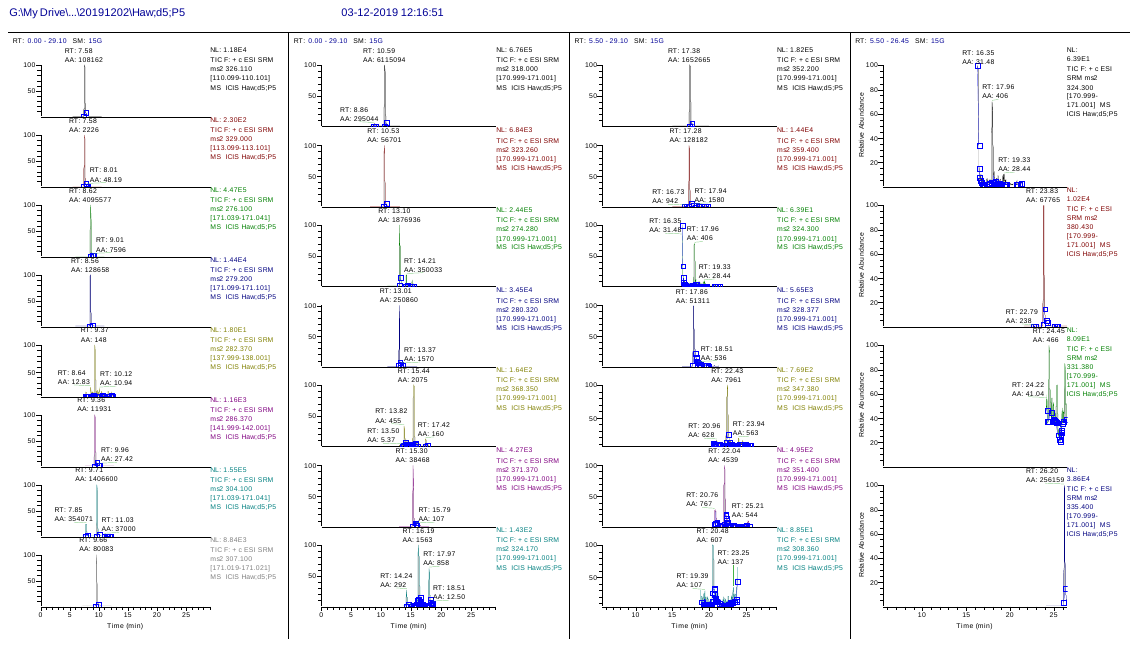

**Figure s6:** **Sample chromatograms obtained from LCMS neurochemical analysis.**

A) Blank (2% ACN prepared in 0.5% FA), B) Standard 12;5 from calibration curve and C) Pooled Hawthorn brain sample. For each neurochemical compound, the molecular weight and retention time details mention in table s11. Note that the blank chromatograms (A) do not show any peak at respected retention times. In contrast, calibration and sample chromatogram indicate the internal standard and compound peaks at particular retention time points where they are eluting.

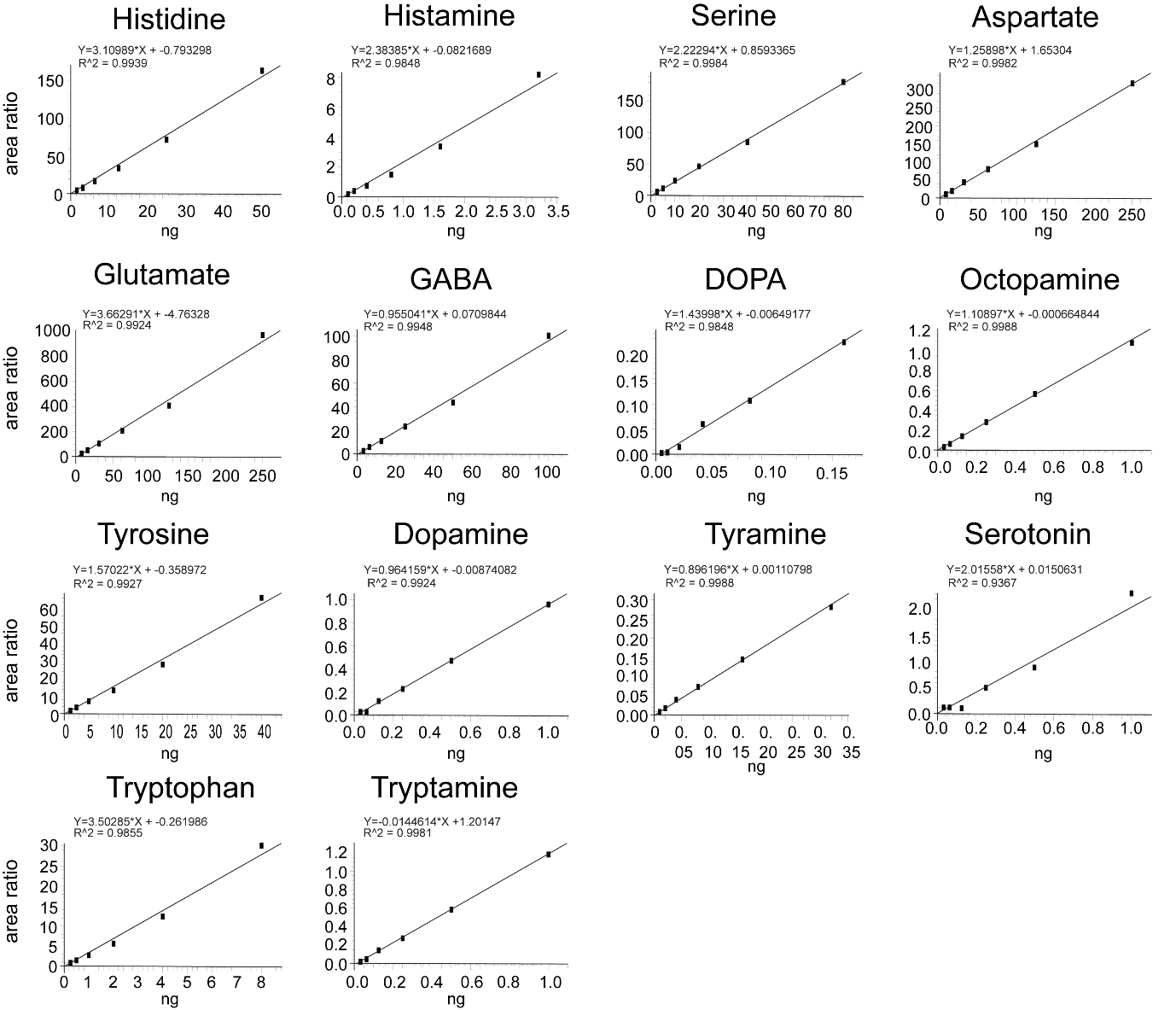

**Figure s7: Sample** **calibration curves for one of the LC-MS runs showing all 14 neurotransmitters analysed.**

**Table s1:** **Total number of pupae sampled after winter in the apple race**

| Total no. of | D 0 | D 5 | D 10 | D 15 | D 20 | D 25 | D 30 | D 35 | D 40 | D 45 | D 50 | D 55 | D 60 | D 65 | D 70 | total |
| --- | --- | --- | --- | --- | --- | --- | --- | --- | --- | --- | --- | --- | --- | --- | --- | --- |
| pupae opened | 27 | 46 | 46 | 70 | 65 | 60 | 37 | 37 | 10 | 0 | 6 | 0 | 3 | 0 | 0 | 407 |
| larval brain,stage3 | 9 | 24 | 30 | 20 | 24 | 11 | 11 | 5 | 0 | 0 | 0 | 0 | 0 | 0 | 0 | 134 |
| adult brain, stage 4 | 11 | 1 | 1 | 3 | 6 | 20 | 13 | 11 | 4 | 0 | 0 | 0 | 0 | 0 | 0 | 70 |
| adult brain, stage 5 | 3 | 8 | 2 | 2 | 4 | 4 | 8 | 6 | 3 | 0 | 3 | 0 | 0 | 0 | 0 | 43 |
| adult brain, stage 6 | 0 | 4 | 5 | 6 | 1 | 1 | 1 | 1 | 0 | 0 | 0 | 0 | 1 | 0 | 0 | 20 |
| dead fly pupae | 3 | 5 | 8 | 38 | 29 | 21 | 0 | 10 | 3 | 0 | 0 | 0 | 0 | 0 | 0 | 117 |
| dead fly | 0 | 0 | 0 | 0 | 0 | 0 | 3 | 1 | 0 | 0 | 3 | 0 | 1 | 0 | 0 | 8 |
| wasp larvae | 0 | 2 | 0 | 1 | 1 | 3 | 0 | 2 | 0 | 0 | 0 | 0 | 1 | 0 | 0 | 10 |
| adult wasp | 1 | 2 | 0 | 0 | 0 | 0 | 1 | 1 | 0 | 0 | 0 | 0 | 0 | 0 | 0 | 5 |

**Table s2:** **Total number of pupae sampled after winter in the hawthorn race**

| Total no. of | D 0 | D 5 | D 10 | D 15 | D 20 | D 25 | D 30 | D 35 | D 40 | D 45 | D 50 | D 55 | D 60 | D 65 | D 70 | total |
| --- | --- | --- | --- | --- | --- | --- | --- | --- | --- | --- | --- | --- | --- | --- | --- | --- |
| pupae opened | 26 | 28 | 56 | 10 | 9 | 50 | 41 | 30 | 55 | 78 | 74 | 31 | 51 | 47 | 19 | 605 |
| larval brain,stage3 | 22 | 25 | 25 | 2 | 6 | 20 | 20 | 19 | 14 | 14 | 2 | 1 | 0 | 1 | 0 | 171 |
| adult brain, stage 4 | 1 | 0 | 0 | 0 | 0 | 0 | 0 | 1 | 13 | 13 | 15 | 4 | 8 | 2 | 3 | 60 |
| adult brain, stage 5 | 0 | 0 | 1 | 0 | 0 | 0 | 0 | 0 | 0 | 3 | 3 | 1 | 16 | 8 | 3 | 35 |
| adult brain, stage 6 | 0 | 0 | 1 | 0 | 0 | 0 | 0 | 0 | 0 | 1 | 4 | 5 | 10 | 15 | 5 | 41 |
| dead fly pupae | 0 | 1 | 0 | 0 | 0 | 2 | 3 | 2 | 14 | 18 | 16 | 4 | 1 | 8 | 1 | 70 |
| dead fly | 0 | 0 | 3 | 0 | 0 | 6 | 2 | 1 | 3 | 9 | 4 | 1 | 5 | 3 | 4 | 41 |
| wasp larvae | 2 | 2 | 15 | 1 | 0 | 9 | 7 | 3 | 4 | 14 | 16 | 7 | 6 | 10 | 3 | 99 |
| adult wasp | 1 | 0 | 11 | 7 | 3 | 13 | 9 | 4 | 7 | 6 | 14 | 8 | 5 | 0 | 0 | 88 |

**Table s3-s6**: Generalised linear mixed models, Tukey’s HSD tests was performed for pairwise multiple comparisons with a significance cut off of 0.0001***, 0.001**, or 0.05* as indicated.

**Table s3: Within and between host race metabolic rate comparisons.**

| compariosns | estimate | SE | df | t.ratio | p.value | |
| --- | --- | --- | --- | --- | --- | --- |
| Apple day 8 - Apple day 20 | -0.0013 | 0.0004 | 237.8199 | -3.1127 | 0.0111 | * |
| Apple day 20 - Hawthorn day 20 | 0.0016 | 0.0005 | 136.0076 | 3.5224 | 0.0032 | ** |
| Apple day 8 - Hawthorn day 8 | 0.0002 | 0.0004 | 75.1650 | 0.5492 | 0.9465 |  |
| Hawthorn day 8 - Hawthorn day 20 | -0.0027 | 0.0004 | 784.8679 | -6.2869 | 0.0001 | *** |

**Table s4: Between host race stage comparisons for each neurotransmitter.**

| Histidine | estimate | SE | df | t.ratio | p.value | |
| --- | --- | --- | --- | --- | --- | --- |
| Apple stage 1 - Hawthorn stage 1 | -1.3238 | 3.4071 | 151.0829 | -0.3885 | 1.0000 |  |
| Apple stage 2 - Hawthorn stage 2 | -4.2289 | 3.3472 | 150.2634 | -1.2634 | 0.9823 |  |
| Apple stage 3 - Hawthorn stage 3 | 2.7567 | 2.2972 | 152.9702 | 1.2000 | 0.9883 |  |
| Apple stage 4 - Hawthorn stage 4 | -9.1510 | 3.5061 | 154.1119 | -2.6100 | 0.2835 |  |
| Apple stage 5 - Hawthorn stage 5 | -1.3582 | 4.7046 | 153.6632 | -0.2887 | 1.0000 |  |
| Apple stage 6 - Hawthorn stage 6 | -1.4942 | 5.4154 | 157.3576 | -0.2759 | 1.0000 |  |
| Apple stage 7 - Hawthorn stage 7 | -7.1651 | 2.0019 | 34.5272 | -3.5791 | 0.0055 | ** |
| Apple stage 8 - Hawthorn stage 8 | -1.5290 | 2.0606 | 35.3436 | -0.7420 | 0.8794 |  |
| Serine | estimate | SE | df | t.ratio | p.value | |
| Apple stage 1 - Hawthorn stage 1 | -4.3476 | 1.8536 | 138.3411 | -2.3454 | 0.4502 |  |
| Apple stage 2 - Hawthorn stage 2 | -1.8663 | 1.9644 | 140.9179 | -0.9501 | 0.9984 |  |
| Apple stage 3 - Hawthorn stage 3 | 2.0409 | 1.2809 | 139.9661 | 1.5934 | 0.9087 |  |
| Apple stage 4 - Hawthorn stage 4 | -2.2410 | 1.9622 | 141.3912 | -1.1421 | 0.9921 |  |
| Apple stage 5 - Hawthorn stage 5 | -7.3387 | 2.6361 | 141.0016 | -2.7839 | 0.1984 |  |
| Apple stage 6 - Hawthorn stage 6 | 11.6866 | 3.0346 | 146.8927 | 3.8511 | 0.0093 | ** |
| Apple stage 7 - Hawthorn stage 7 | -2.2881 | 2.1287 | 33.5397 | -1.0749 | 0.7069 |  |
| Apple stage 8 - Hawthorn stage 8 | 1.4459 | 2.2274 | 34.2703 | 0.6491 | 0.9151 |  |
| Histamine | estimate | SE | df | t.ratio | p.value | |
| Apple stage 1 - Hawthorn stage 1 | 0.0024 | 0.0052 | 123.6331 | 0.4560 | 1.0000 |  |
| Apple stage 2 - Hawthorn stage 2 | 0.0075 | 0.0048 | 122.3740 | 1.5637 | 0.9186 |  |
| Apple stage 3 - Hawthorn stage 3 | 0.0009 | 0.0032 | 125.6815 | 0.2744 | 1.0000 |  |
| Apple stage 4 - Hawthorn stage 4 | -0.0107 | 0.0043 | 123.2558 | -2.4914 | 0.3553 |  |
| Apple stage 5 - Hawthorn stage 5 | -0.0087 | 0.0058 | 124.1761 | -1.4890 | 0.9409 |  |
| Apple stage 6 - Hawthorn stage 6 | -0.0775 | 0.0067 | 124.0605 | -11.5464 | 0.0001 | *** |
| Apple stage 7 - Hawthorn stage 7 | -0.0231 | 0.0476 | 33.1454 | -0.4861 | 0.9617 |  |
| Apple stage 8 - Hawthorn stage 8 | -0.1543 | 0.0502 | 35.5456 | -3.0755 | 0.0201 | * |
| Aspartate | estimate | SE | df | t.ratio | p.value | |
| Apple stage 1 - Hawthorn stage 1 | -3.6951 | 3.9919 | 150.1434 | -0.9256 | 0.9988 |  |
| Apple stage 2 - Hawthorn stage 2 | -0.8473 | 3.7653 | 148.7253 | -0.2250 | 1.0000 |  |
| Apple stage 3 - Hawthorn stage 3 | 2.8912 | 2.6426 | 149.8157 | 1.0941 | 0.9945 |  |
| Apple stage 4 - Hawthorn stage 4 | -14.2007 | 3.9688 | 150.4097 | -3.5781 | 0.0228 | * |
| Apple stage 5 - Hawthorn stage 5 | -4.4480 | 5.2702 | 149.2740 | -0.8440 | 0.9995 |  |
| Apple stage 6 - Hawthorn stage 6 | -11.8885 | 6.1894 | 150.9730 | -1.9208 | 0.7444 |  |
| Apple stage 7 - Hawthorn stage 7 | -8.6402 | 12.0732 | 34.0117 | -0.7156 | 0.8902 |  |
| Apple stage 8 - Hawthorn stage 8 | 9.1575 | 12.4722 | 34.4754 | 0.7342 | 0.8826 |  |
| Glutamate | estimate | SE | df | t.ratio | p.value | |
| Apple stage 1 - Hawthorn stage 1 | -15.3468 | 7.3757 | 150.9329 | -2.0807 | 0.6371 |  |
| Apple stage 2 - Hawthorn stage 2 | -11.1878 | 7.5039 | 150.0898 | -1.4909 | 0.9410 |  |
| Apple stage 3 - Hawthorn stage 3 | 2.5626 | 4.8625 | 150.9665 | 0.5270 | 1.0000 |  |
| Apple stage 4 - Hawthorn stage 4 | -30.9558 | 7.2703 | 152.3285 | -4.2578 | 0.0021 | ** |
| Apple stage 5 - Hawthorn stage 5 | -5.3290 | 9.5928 | 151.8868 | -0.5555 | 1.0000 |  |
| Apple stage 6 - Hawthorn stage 6 | -24.0413 | 11.5010 | 153.2042 | -2.0904 | 0.6303 |  |
| Apple stage 7 - Hawthorn stage 7 | -15.4941 | 10.3199 | 33.0780 | -1.5014 | 0.4481 |  |
| Apple stage 8 - Hawthorn stage 8 | -21.4532 | 11.4093 | 34.6585 | -1.8803 | 0.2550 |  |
| GABA | estimate | SE | df | t.ratio | p.value | |
| Apple stage 1 - Hawthorn stage 1 | -0.3228 | 0.6631 | 149.8323 | -0.4868 | 1.0000 |  |
| Apple stage 2 - Hawthorn stage 2 | 0.5101 | 0.6595 | 146.8546 | 0.7736 | 0.9998 |  |
| Apple stage 3 - Hawthorn stage 3 | -0.8280 | 0.4311 | 147.5445 | -1.9207 | 0.7444 |  |
| Apple stage 4 - Hawthorn stage 4 | -0.6508 | 0.6661 | 153.4659 | -0.9770 | 0.9980 |  |
| Apple stage 5 - Hawthorn stage 5 | -4.3573 | 0.8868 | 150.7387 | -4.9138 | 0.0001 | *** |
| Apple stage 6 - Hawthorn stage 6 | -13.4738 | 1.1482 | 151.4194 | -11.7348 | 0.0000 | *** |
| Apple stage 7 - Hawthorn stage 7 | -3.4803 | 3.1175 | 32.8182 | -1.1164 | 0.6822 |  |
| Apple stage 8 - Hawthorn stage 8 | -0.4529 | 3.4660 | 34.2058 | -0.1307 | 0.9992 |  |
| Dopa | estimate | SE | df | t.ratio | p.value | |
| Apple stage 1 - Hawthorn stage 1 | -0.0312 | 0.0264 | 149.9665 | -1.1818 | 0.9896 |  |
| Apple stage 2 - Hawthorn stage 2 | 0.0066 | 0.0273 | 151.0489 | 0.2408 | 1.0000 |  |
| Apple stage 3 - Hawthorn stage 3 | 0.0361 | 0.0180 | 150.3388 | 2.0058 | 0.6888 |  |
| Apple stage 4 - Hawthorn stage 4 | -0.1378 | 0.0295 | 151.2967 | -4.6798 | 0.0004 | *** |
| Apple stage 5 - Hawthorn stage 5 | 0.0591 | 0.0368 | 151.8817 | 1.6048 | 0.9048 |  |
| Apple stage 6 - Hawthorn stage 6 | 0.0076 | 0.0477 | 153.2333 | 0.1594 | 1.0000 |  |
| Apple stage 7 - Hawthorn stage 7 | -0.0009 | 0.0017 | 32.4581 | -0.5609 | 0.9429 |  |
| Apple stage 8 - Hawthorn stage 8 | 0.0014 | 0.0019 | 33.3627 | 0.7705 | 0.8672 |  |
| Octopamine | estimate | SE | df | t.ratio | p.value | |
| Apple stage 1 - Hawthorn stage 1 | -0.0006 | 0.0476 | 76.5057 | -0.0126 | 1.0000 |  |
| Apple stage 2 - Hawthorn stage 2 | 0.0000 | 0.0447 | 78.2365 | 0.0003 | 1.0000 |  |
| Apple stage 3 - Hawthorn stage 3 | 0.0255 | 0.0349 | 83.5626 | 0.7329 | 0.9999 |  |
| Apple stage 4 - Hawthorn stage 4 | 0.0562 | 0.0333 | 80.0402 | 1.6869 | 0.8688 |  |
| Apple stage 5 - Hawthorn stage 5 | 0.1658 | 0.0452 | 83.0806 | 3.6674 | 0.0206 | * |
| Apple stage 6 - Hawthorn stage 6 | -0.0460 | 0.0511 | 81.5941 | -0.8994 | 0.9990 |  |
| Apple stage 7 - Hawthorn stage 7 | -0.0331 | 0.0272 | 32.5550 | -1.2151 | 0.6219 |  |
| Apple stage 8 - Hawthorn stage 8 | -0.1006 | 0.0304 | 34.5325 | -3.3061 | 0.0113 | * |
| Tyrosine | estimate | SE | df | t.ratio | p.value | |
| Apple stage 1 - Hawthorn stage 1 | -0.8273 | 3.1443 | 143.1745 | -0.2631 | 1.0000 |  |
| Apple stage 2 - Hawthorn stage 2 | -0.4517 | 3.5577 | 144.4147 | -0.1270 | 1.0000 |  |
| Apple stage 3 - Hawthorn stage 3 | 2.0068 | 2.2841 | 150.3338 | 0.8786 | 0.9992 |  |
| Apple stage 4 - Hawthorn stage 4 | -29.1204 | 3.3965 | 151.2312 | -8.5737 | 0.0001 | *** |
| Apple stage 5 - Hawthorn stage 5 | -13.2214 | 4.4727 | 145.9859 | -2.9560 | 0.1329 |  |
| Apple stage 6 - Hawthorn stage 6 | -18.4293 | 5.2246 | 152.3163 | -3.5274 | 0.0267 |  |
| Apple stage 7 - Hawthorn stage 7 | -1.3088 | 1.0043 | 33.5635 | -1.3032 | 0.5674 |  |
| Apple stage 8 - Hawthorn stage 8 | 0.6938 | 1.0493 | 34.4397 | 0.6612 | 0.9109 |  |
| Dopamine | estimate | SE | df | t.ratio | p.value | |
| Apple stage 1 - Hawthorn stage 1 | 0.0095 | 0.0063 | 128.4363 | 1.5266 | 0.9304 |  |
| Apple stage 2 - Hawthorn stage 2 | -0.0112 | 0.0067 | 131.8681 | -1.6715 | 0.8778 |  |
| Apple stage 3 - Hawthorn stage 3 | 0.0030 | 0.0053 | 138.6630 | 0.5725 | 1.0000 |  |
| Apple stage 4 - Hawthorn stage 4 | 0.0022 | 0.0069 | 131.5225 | 0.3136 | 1.0000 |  |
| Apple stage 5 - Hawthorn stage 5 | 0.0440 | 0.0089 | 134.1271 | 4.9272 | 0.0001 | *** |
| Apple stage 6 - Hawthorn stage 6 | 0.0211 | 0.0108 | 135.5962 | 1.9635 | 0.7168 |  |
| Apple stage 7 - Hawthorn stage 7 | -0.0829 | 0.0529 | 31.2879 | -1.5692 | 0.4102 |  |
| Apple stage 8 - Hawthorn stage 8 | 0.0934 | 0.0512 | 31.1948 | 1.8223 | 0.2823 |  |
| Tyramine | estimate | SE | df | t.ratio | p.value | |
| Apple stage 1 - Hawthorn stage 1 | 0.0018 | 0.0031 | 137.0627 | 0.5680 | 1.0000 |  |
| Apple stage 2 - Hawthorn stage 2 | 0.0045 | 0.0029 | 134.6750 | 1.5361 | 0.9277 |  |
| Apple stage 3 - Hawthorn stage 3 | 0.0009 | 0.0020 | 135.1031 | 0.4848 | 1.0000 |  |
| Apple stage 4 - Hawthorn stage 4 | -0.0173 | 0.0028 | 136.4020 | -6.1011 | 0.0001 | *** |
| Apple stage 5 - Hawthorn stage 5 | -0.0068 | 0.0037 | 136.8321 | -1.8537 | 0.7850 |  |
| Apple stage 6 - Hawthorn stage 6 | -0.0154 | 0.0045 | 137.4406 | -3.4070 | 0.0393 | * |
| Apple stage 7 - Hawthorn stage 7 | 0.0127 | 0.0110 | 31.2861 | 1.1570 | 0.6577 |  |
| Apple stage 8 - Hawthorn stage 8 | 0.0358 | 0.0108 | 32.0899 | 3.3242 | 0.0114 | * |
| Serotonine | estimate | SE | df | t.ratio | p.value | |
| Apple stage 1 - Hawthorn stage 1 | 0.0009 | 0.0279 | 89.5853 | 0.0329 | 1.0000 |  |
| Apple stage 2 - Hawthorn stage 2 | 0.0085 | 0.0291 | 85.6962 | 0.2903 | 1.0000 |  |
| Apple stage 3 - Hawthorn stage 3 | -0.0133 | 0.0217 | 87.6624 | -0.6126 | 1.0000 |  |
| Apple stage 4 - Hawthorn stage 4 | 0.0466 | 0.0416 | 94.9574 | 1.1204 | 0.9930 |  |
| Apple stage 5 - Hawthorn stage 5 | 0.0660 | 0.0458 | 88.5421 | 1.4405 | 0.9519 |  |
| Apple stage 6 - Hawthorn stage 6 | 0.4526 | 0.0462 | 91.0937 | 9.7915 | 0.0001 | *** |
| Apple stage 7 - Hawthorn stage 7 | -0.1029 | 0.1705 | 24.5415 | -0.6036 | 0.9299 |  |
| Apple stage 8 - Hawthorn stage 8 | 0.3304 | 0.1702 | 26.8478 | 1.9415 | 0.2352 |  |
| Tryptophan | estimate | SE | df | t.ratio | p.value | |
| Apple stage 1 - Hawthorn stage 1 | -0.1678 | 0.3689 | 153.5635 | -0.4549 | 1.0000 |  |
| Apple stage 2 - Hawthorn stage 2 | -0.0997 | 0.3661 | 153.1281 | -0.2724 | 1.0000 |  |
| Apple stage 3 - Hawthorn stage 3 | 0.4235 | 0.2410 | 153.2847 | 1.7576 | 0.8378 |  |
| Apple stage 4 - Hawthorn stage 4 | -1.5851 | 0.3798 | 153.9048 | -4.1734 | 0.0028 | ** |
| Apple stage 5 - Hawthorn stage 5 | 0.0957 | 0.4913 | 153.9227 | 0.1947 | 1.0000 |  |
| Apple stage 6 - Hawthorn stage 6 | 1.1234 | 0.6015 | 154.2040 | 1.8678 | 0.7770 |  |
| Apple stage 7 - Hawthorn stage 7 | -1.0653 | 0.6399 | 22.0000 | -1.6650 | 0.3651 |  |
| Apple stage 8 - Hawthorn stage 8 | 1.5244 | 0.6623 | 22.0000 | 2.3016 | 0.1283 |  |
| Tryptamine | estimate | SE | df | t.ratio | p.value | |
| Apple stage 1 - Hawthorn stage 1 | -0.0155 | 0.0128 | 117.2359 | -1.2105 | 0.9872 |  |
| Apple stage 2 - Hawthorn stage 2 | 0.0075 | 0.0116 | 116.3439 | 0.6443 | 1.0000 |  |
| Apple stage 3 - Hawthorn stage 3 | 0.0004 | 0.0078 | 115.6457 | 0.0478 | 1.0000 |  |
| Apple stage 4 - Hawthorn stage 4 | -0.0045 | 0.0097 | 117.6350 | -0.4698 | 1.0000 |  |
| Apple stage 5 - Hawthorn stage 5 | 0.0040 | 0.0136 | 115.5865 | 0.2917 | 1.0000 |  |
| Apple stage 6 - Hawthorn stage 6 | 0.0075 | 0.0154 | 118.8689 | 0.4883 | 1.0000 |  |
| Apple stage 7 - Hawthorn stage 7 | -0.0566 | 0.0167 | 33.8268 | -3.3831 | 0.0094 | ** |
| Apple stage 8 - Hawthorn stage 8 | 0.0235 | 0.0183 | 35.9997 | 1.2844 | 0.5786 |  |

**Table s5: Apple race developmental stage comparisons for each neurotransmitter.**

| Histidine | estimate | SE | df | t.ratio | p.value | |
| --- | --- | --- | --- | --- | --- | --- |
| Stage 1 - stage 2 | 6.3710 | 3.4071 | 151.0829 | 1.8699 | 0.7757 |  |
| Stage 1 - stage 3 | -4.0614 | 3.5846 | 33.8738 | -1.1330 | 0.9907 |  |
| Stage 1 - stage 4 | -10.7407 | 3.8992 | 47.0710 | -2.7546 | 0.2328 |  |
| Stage 1 - stage 5 | 1.3674 | 4.3923 | 66.0851 | 0.3113 | 1.0000 |  |
| Stage 1 - stage 6 | -4.7250 | 5.4277 | 104.0757 | -0.8705 | 0.9993 |  |
| Stage 2 - stage 3 | 10.4324 | 3.7043 | 38.2197 | 2.8163 | 0.2135 |  |
| Stage 2 - stage 4 | -4.3698 | 4.0095 | 51.7618 | -1.0898 | 0.9939 |  |
| Stage 2 - stage 5 | 5.0036 | 4.4905 | 70.5918 | 1.1143 | 0.9930 |  |
| Stage 2 - stage 6 | 11.0960 | 5.5075 | 107.2170 | 2.0147 | 0.6825 |  |
| Stage 3 - stage 4 | -14.8022 | 3.0304 | 161.9721 | -4.8845 | 0.0002 | *** |
| Stage 3 - stage 5 | 5.4288 | 3.6700 | 161.9028 | 1.4792 | 0.9442 |  |
| Stage 3 - stage 6 | -0.6636 | 4.8474 | 161.6184 | -0.1369 | 1.0000 |  |
| Stage 4 - stage 5 | -9.3734 | 3.8692 | 154.6094 | -2.4226 | 0.3975 |  |
| Stage 4 - stage 6 | -15.4657 | 5.0646 | 158.8954 | -3.0537 | 0.1032 |  |
| Stage 5 - stage 6 | -6.0924 | 5.3788 | 154.4870 | -1.1327 | 0.9927 |  |
| Stage 6 - stage 7 | 1.2020 | 5.4286 | 126.1111 | 0.2214 | 1.0000 |  |
| Stage 7 - stage 8 | -3.8320 | 2.0147 | 34.3189 | -1.9020 | 0.2461 |  |
| Serine | estimate | SE | df | t.ratio | p.value | |
| Stage 1 - stage 2 | 1.0629 | 1.9492 | 138.0375 | 0.5453 | 1.0000 |  |
| Stage 1 - stage 3 | 1.0320 | 2.0756 | 22.0955 | 0.4972 | 1.0000 |  |
| Stage 1 - stage 4 | -12.1406 | 2.2253 | 30.0121 | -5.4557 | 0.0003 | *** |
| Stage 1 - stage 5 | 10.0534 | 2.5654 | 47.8072 | 3.9189 | 0.0134 | * |
| Stage 1 - stage 6 | 16.2475 | 3.0339 | 76.3100 | 5.3553 | 0.0001 | *** |
| Stage 2 - stage 3 | 0.0309 | 2.2305 | 28.7229 | 0.0139 | 1.0000 |  |
| Stage 2 - stage 4 | -11.0777 | 2.3705 | 37.3524 | -4.6732 | 0.0020 | ** |
| Stage 2 - stage 5 | -8.9904 | 2.6922 | 55.3685 | -3.3394 | 0.0602 |  |
| Stage 2 - stage 6 | -15.1846 | 3.1419 | 82.7951 | -4.8329 | 0.0004 | *** |
| Stage 3 - stage 4 | -11.1086 | 1.6765 | 152.9994 | -6.6259 | 0.0001 | *** |
| Stage 3 - stage 5 | 9.0214 | 2.1364 | 152.8918 | 4.2227 | 0.0024 | ** |
| Stage 3 - stage 6 | 15.2155 | 2.6621 | 151.7478 | 5.7157 | 0.0001 | *** |
| Stage 4 - stage 5 | -2.0872 | 2.2173 | 144.3248 | -0.9413 | 0.9985 |  |
| Stage 4 - stage 6 | 4.1069 | 2.7698 | 147.4602 | 1.4828 | 0.9431 |  |
| Stage 5 - stage 6 | 6.1941 | 2.9927 | 141.4463 | 2.0697 | 0.6448 |  |
| Stage 6 - stage 7 | -5.0133 | 3.3908 | 83.9981 | -1.4785 | 0.9821 |  |
| Stage 7 - stage 8 | -1.9049 | 2.1402 | 33.2748 | -0.8900 | 0.8100 |  |
| Histamine | estimate | SE | df | t.ratio | p.value | |
| Stage 1 - stage 2 | 0.0081 | 0.0054 | 124.0529 | 1.4793 | 0.9435 |  |
| Stage 1 - stage 3 | 0.0044 | 0.0073 | 22.0036 | 0.5997 | 1.0000 |  |
| Stage 1 - stage 4 | -0.0114 | 0.0076 | 25.0608 | -1.5019 | 0.9265 |  |
| Stage 1 - stage 5 | 0.0120 | 0.0080 | 31.2531 | 1.4888 | 0.9328 |  |
| Stage 1 - stage 6 | 0.0187 | 0.0088 | 43.3008 | 2.1158 | 0.6149 |  |
| Stage 2 - stage 3 | 0.0037 | 0.0069 | 18.5033 | 0.5353 | 1.0000 |  |
| Stage 2 - stage 4 | -0.0033 | 0.0072 | 21.4419 | -0.4550 | 1.0000 |  |
| Stage 2 - stage 5 | -0.0039 | 0.0077 | 27.4179 | -0.5041 | 1.0000 |  |
| Stage 2 - stage 6 | -0.0107 | 0.0086 | 39.3271 | -1.2439 | 0.9814 |  |
| Stage 3 - stage 4 | -0.0070 | 0.0040 | 128.5870 | -1.7647 | 0.8337 |  |
| Stage 3 - stage 5 | 0.0076 | 0.0048 | 126.9465 | 1.5858 | 0.9111 |  |
| Stage 3 - stage 6 | 0.0144 | 0.0061 | 126.4946 | 2.3716 | 0.4329 |  |
| Stage 4 - stage 5 | 0.0006 | 0.0050 | 123.0837 | 0.1194 | 1.0000 |  |
| Stage 4 - stage 6 | 0.0074 | 0.0063 | 124.2264 | 1.1637 | 0.9907 |  |
| Stage 5 - stage 6 | 0.0068 | 0.0068 | 122.5118 | 1.0012 | 0.9974 |  |
| Stage 6 - stage 7 | 0.5474 | 0.0367 | 65.1478 | 14.9044 | 0.0001 | *** |
| Stage 7 - stage 8 | 0.1078 | 0.0490 | 32.2913 | 2.1992 | 0.1450 |  |
| Aspartate | estimate | SE | df | t.ratio | p.value | |
| Stage 1 - stage 2 | 1.3990 | 3.9919 | 150.1434 | 0.3505 | 1.0000 |  |
| Stage 1 - stage 3 | 9.2458 | 7.3079 | 14.8824 | 1.2652 | 0.9720 |  |
| Stage 1 - stage 4 | -40.6847 | 7.4747 | 16.3835 | -5.4430 | 0.0021 | ** |
| Stage 1 - stage 5 | 29.2739 | 7.9529 | 20.6859 | 3.6809 | 0.0481 | * |
| Stage 1 - stage 6 | 17.3012 | 8.6076 | 28.0209 | 2.0100 | 0.6836 |  |
| Stage 2 - stage 3 | -7.8468 | 7.2160 | 14.2074 | -1.0874 | 0.9904 |  |
| Stage 2 - stage 4 | -39.2857 | 7.3849 | 15.6796 | -5.3198 | 0.0029 | ** |
| Stage 2 - stage 5 | -27.8749 | 7.8686 | 19.9217 | -3.5426 | 0.0655 |  |
| Stage 2 - stage 6 | -15.9022 | 8.5297 | 27.1797 | -1.8643 | 0.7697 |  |
| Stage 3 - stage 4 | -31.4389 | 3.7533 | 156.0706 | -8.3763 | 0.0001 | *** |
| Stage 3 - stage 5 | 20.0281 | 4.7937 | 158.0994 | 4.1780 | 0.0028 | ** |
| Stage 3 - stage 6 | 8.0554 | 5.6950 | 154.3405 | 1.4145 | 0.9591 |  |
| Stage 4 - stage 5 | -11.4108 | 4.6537 | 150.4490 | -2.4520 | 0.3785 |  |
| Stage 4 - stage 6 | -23.3835 | 5.8111 | 150.4609 | -4.0239 | 0.0050 | ** |
| Stage 5 - stage 6 | -11.9727 | 6.2388 | 149.3851 | -1.9191 | 0.7455 |  |
| Stage 6 - stage 7 | 140.8102 | 15.4588 | 28.1886 | 9.1088 | 0.0001 | *** |
| Stage 7 - stage 8 | 19.7866 | 12.1420 | 33.9396 | 1.6296 | 0.3761 |  |
| Glutamate | estimate | SE | df | t.ratio | p.value | |
| Stage 1 - stage 2 | 14.4270 | 7.6977 | 151.1464 | 1.8742 | 0.7731 |  |
| Stage 1 - stage 3 | 8.1177 | 9.8073 | 23.4441 | 0.8277 | 0.9993 |  |
| Stage 1 - stage 4 | -38.4449 | 10.2692 | 28.3157 | -3.7437 | 0.0321 | * |
| Stage 1 - stage 5 | 5.2307 | 11.1385 | 37.5008 | 0.4696 | 1.0000 |  |
| Stage 1 - stage 6 | -14.6877 | 13.0425 | 61.9126 | -1.1261 | 0.9922 |  |
| Stage 2 - stage 3 | 6.3093 | 9.9010 | 24.4487 | 0.6372 | 0.9999 |  |
| Stage 2 - stage 4 | -24.0178 | 10.3587 | 29.4102 | -2.3186 | 0.4864 |  |
| Stage 2 - stage 5 | 9.1963 | 11.2211 | 38.6990 | 0.8196 | 0.9995 |  |
| Stage 2 - stage 6 | 29.1148 | 13.1131 | 63.2049 | 2.2203 | 0.5418 |  |
| Stage 3 - stage 4 | -30.3272 | 6.6180 | 159.0077 | -4.5825 | 0.0006 | *** |
| Stage 3 - stage 5 | -2.8870 | 7.9892 | 159.6720 | -0.3614 | 1.0000 |  |
| Stage 3 - stage 6 | -22.8054 | 10.4302 | 157.5626 | -2.1865 | 0.5619 |  |
| Stage 4 - stage 5 | -33.2141 | 8.1954 | 151.8280 | -4.0528 | 0.0045 | ** |
| Stage 4 - stage 6 | -53.1326 | 10.7735 | 153.8021 | -4.9318 | 0.0001 | *** |
| Stage 5 - stage 6 | -19.9185 | 11.3870 | 151.1367 | -1.7492 | 0.8420 |  |
| Stage 6 - stage 7 | 138.7044 | 14.7222 | 70.0816 | 9.4215 | 0.0001 | *** |
| Stage 7 - stage 8 | -5.3597 | 10.4487 | 33.5306 | -0.5130 | 0.9554 |  |
| GABA | estimate | SE | df | t.ratio | p.value | |
| Stage 1 - stage 2 | 1.2415 | 0.6770 | 150.3439 | 1.8337 | 0.7968 |  |
| Stage 1 - stage 3 | -0.6305 | 0.6577 | 48.2274 | -0.9586 | 0.9979 |  |
| Stage 1 - stage 4 | 2.4515 | 0.7364 | 69.1423 | 3.3288 | 0.0577 | * |
| Stage 1 - stage 5 | 0.5087 | 0.8492 | 95.4747 | 0.5990 | 1.0000 |  |
| Stage 1 - stage 6 | 0.9161 | 1.1295 | 130.6834 | 0.8111 | 0.9996 |  |
| Stage 2 - stage 3 | 1.8720 | 0.6565 | 48.5321 | 2.8514 | 0.1915 |  |
| Stage 2 - stage 4 | 3.6929 | 0.7354 | 69.5930 | 5.0219 | 0.0002 | *** |
| Stage 2 - stage 5 | 0.7328 | 0.8482 | 96.0020 | 0.8639 | 0.9993 |  |
| Stage 2 - stage 6 | 0.3254 | 1.1288 | 131.0409 | 0.2883 | 1.0000 |  |
| Stage 3 - stage 4 | 1.8209 | 0.5774 | 157.9992 | 3.1534 | 0.0793 |  |
| Stage 3 - stage 5 | 1.1392 | 0.7232 | 157.5404 | 1.5752 | 0.9155 |  |
| Stage 3 - stage 6 | 1.5466 | 1.0421 | 157.3267 | 1.4842 | 0.9429 |  |
| Stage 4 - stage 5 | 2.9601 | 0.7860 | 155.4472 | 3.7659 | 0.0122 | * |
| Stage 4 - stage 6 | 3.3675 | 1.0924 | 157.9889 | 3.0826 | 0.0958 |  |
| Stage 5 - stage 6 | 0.4074 | 1.1487 | 151.7796 | 0.3547 | 1.0000 |  |
| Stage 6 - stage 7 | 54.6217 | 3.1419 | 48.2363 | 17.3849 | 0.0001 | *** |
| Stage 7 - stage 8 | 3.1468 | 3.1622 | 33.4048 | 0.9951 | 0.7532 |  |
| Dopa | estimate | SE | df | t.ratio | p.value | |
| Stage 1 - stage 2 | 0.0226 | 0.0277 | 151.2025 | 0.8146 | 0.9996 |  |
| Stage 1 - stage 3 | 0.0100 | 0.0262 | 51.4381 | 0.3824 | 1.0000 |  |
| Stage 1 - stage 4 | -0.0903 | 0.0300 | 76.8582 | -3.0122 | 0.1250 |  |
| Stage 1 - stage 5 | 0.0839 | 0.0344 | 99.0306 | 2.4369 | 0.3920 |  |
| Stage 1 - stage 6 | 0.0381 | 0.0461 | 135.8433 | 0.8257 | 0.9996 |  |
| Stage 2 - stage 3 | 0.0126 | 0.0280 | 61.1593 | 0.4493 | 1.0000 |  |
| Stage 2 - stage 4 | -0.0677 | 0.0316 | 85.0699 | -2.1440 | 0.5935 |  |
| Stage 2 - stage 5 | -0.0613 | 0.0358 | 104.8080 | -1.7109 | 0.8592 |  |
| Stage 2 - stage 6 | -0.0155 | 0.0472 | 137.5124 | -0.3286 | 1.0000 |  |
| Stage 3 - stage 4 | -0.0803 | 0.0250 | 156.1366 | -3.2082 | 0.0684 |  |
| Stage 3 - stage 5 | 0.0739 | 0.0305 | 154.7410 | 2.4243 | 0.3964 |  |
| Stage 3 - stage 6 | 0.0281 | 0.0433 | 157.5659 | 0.6487 | 1.0000 |  |
| Stage 4 - stage 5 | -0.0064 | 0.0328 | 152.0740 | -0.1953 | 1.0000 |  |
| Stage 4 - stage 6 | -0.0522 | 0.0454 | 157.3919 | -1.1481 | 0.9919 |  |
| Stage 5 - stage 6 | -0.0458 | 0.0478 | 153.4026 | -0.9575 | 0.9983 |  |
| Stage 6 - stage 7 | -0.0595 | 0.0438 | 164.5738 | -1.3589 | 0.9928 |  |
| Stage 7 - stage 8 | -0.0036 | 0.0017 | 32.3078 | -2.1342 | 0.1639 |  |
| Octopamine | estimate | SE | df | t.ratio | p.value | |
| Stage 1 - stage 2 | 0.0012 | 0.0476 | 76.5057 | 0.0252 | 1.0000 |  |
| Stage 1 - stage 3 | 0.0400 | 0.0579 | 14.2248 | 0.6908 | 0.9998 |  |
| Stage 1 - stage 4 | -0.0636 | 0.0579 | 14.3025 | -1.0975 | 0.9897 |  |
| Stage 1 - stage 5 | 0.1777 | 0.0600 | 16.7666 | 2.9586 | 0.2045 |  |
| Stage 1 - stage 6 | 0.0462 | 0.0671 | 23.8007 | 0.6889 | 0.9999 |  |
| Stage 2 - stage 3 | -0.0388 | 0.0579 | 14.2248 | -0.6700 | 0.9999 |  |
| Stage 2 - stage 4 | -0.0624 | 0.0579 | 14.3025 | -1.0768 | 0.9911 |  |
| Stage 2 - stage 5 | -0.1765 | 0.0600 | 16.7666 | -2.9386 | 0.2111 |  |
| Stage 2 - stage 6 | -0.0450 | 0.0671 | 23.8007 | -0.6710 | 0.9999 |  |
| Stage 3 - stage 4 | -0.0236 | 0.0303 | 79.8946 | -0.7775 | 0.9997 |  |
| Stage 3 - stage 5 | 0.1376 | 0.0366 | 88.3850 | 3.7659 | 0.0148 | * |
| Stage 3 - stage 6 | 0.0062 | 0.0463 | 86.0316 | 0.1341 | 1.0000 |  |
| Stage 4 - stage 5 | 0.1141 | 0.0355 | 85.9109 | 3.2161 | 0.0737 |  |
| Stage 4 - stage 6 | -0.0174 | 0.0468 | 85.3827 | -0.3711 | 1.0000 |  |
| Stage 5 - stage 6 | -0.1314 | 0.0482 | 82.6230 | -2.7294 | 0.2309 |  |
| Stage 6 - stage 7 | 0.2658 | 0.0531 | 52.0130 | 5.0052 | 0.0007 | *** |
| Stage 7 - stage 8 | 0.0347 | 0.0275 | 31.0306 | 1.2607 | 0.5941 |  |
| Tyrosine | estimate | SE | df | t.ratio | p.value | |
| Stage 1 - stage 2 | 0.2142 | 3.5059 | 143.8441 | 0.0611 | 1.0000 |  |
| Stage 1 - stage 3 | 3.7268 | 3.7433 | 20.9495 | 0.9956 | 0.9961 |  |
| Stage 1 - stage 4 | -31.5498 | 3.9515 | 27.6299 | -7.9844 | 0.0001 | *** |
| Stage 1 - stage 5 | 29.0889 | 4.5641 | 43.7012 | 6.3734 | 0.0001 | *** |
| Stage 1 - stage 6 | 14.4145 | 5.3614 | 72.4407 | 2.6886 | 0.2531 |  |
| Stage 2 - stage 3 | -3.5126 | 4.1441 | 30.7036 | -0.8476 | 0.9992 |  |
| Stage 2 - stage 4 | -31.3357 | 4.3330 | 38.5610 | -7.2319 | 0.0001 | *** |
| Stage 2 - stage 5 | -28.8748 | 4.8982 | 55.0465 | -5.8950 | 0.0001 | *** |
| Stage 2 - stage 6 | -14.2003 | 5.6485 | 82.9349 | -2.5140 | 0.3463 |  |
| Stage 3 - stage 4 | -27.8231 | 3.1299 | 154.2706 | -8.8894 | 0.0001 | *** |
| Stage 3 - stage 5 | 25.3622 | 3.9646 | 149.9901 | 6.3971 | 0.0001 | *** |
| Stage 3 - stage 6 | 10.6877 | 4.7689 | 158.9345 | 2.2411 | 0.5228 |  |
| Stage 4 - stage 5 | -2.4609 | 3.9085 | 150.9554 | -0.6296 | 1.0000 |  |
| Stage 4 - stage 6 | -17.1354 | 4.9327 | 154.8341 | -3.4739 | 0.0314 | * |
| Stage 5 - stage 6 | -14.6745 | 5.2982 | 148.2074 | -2.7697 | 0.2041 |  |
| Stage 6 - stage 7 | -8.1667 | 5.1786 | 105.7313 | -1.5770 | 0.9691 |  |
| Stage 7 - stage 8 | -3.0430 | 1.0103 | 33.1804 | -3.0120 | 0.0243 | * |
| Dopamine | estimate | SE | df | t.ratio | p.value | |
| Stage 1 - stage 2 | -0.0029 | 0.0066 | 129.5853 | -0.4429 | 1.0000 |  |
| Stage 1 - stage 3 | -0.0184 | 0.0066 | 39.6540 | -2.7751 | 0.2294 |  |
| Stage 1 - stage 4 | 0.0104 | 0.0073 | 52.0272 | 1.4298 | 0.9519 |  |
| Stage 1 - stage 5 | 0.0370 | 0.0081 | 68.5808 | 4.5767 | 0.0012 | ** |
| Stage 1 - stage 6 | 0.0359 | 0.0101 | 98.8083 | 3.5537 | 0.0274 | * |
| Stage 2 - stage 3 | 0.0155 | 0.0070 | 47.0390 | 2.2013 | 0.5569 |  |
| Stage 2 - stage 4 | 0.0075 | 0.0076 | 59.0314 | 0.9793 | 0.9976 |  |
| Stage 2 - stage 5 | -0.0399 | 0.0084 | 74.5799 | -4.7376 | 0.0006 | *** |
| Stage 2 - stage 6 | -0.0388 | 0.0104 | 102.0448 | -3.7402 | 0.0152 | * |
| Stage 3 - stage 4 | -0.0080 | 0.0059 | 138.9777 | -1.3546 | 0.9699 |  |
| Stage 3 - stage 5 | 0.0554 | 0.0070 | 138.9975 | 7.9619 | 0.0001 | *** |
| Stage 3 - stage 6 | 0.0543 | 0.0092 | 138.9965 | 5.9004 | 0.0001 | *** |
| Stage 4 - stage 5 | 0.0474 | 0.0074 | 132.6073 | 6.4371 | 0.0001 | *** |
| Stage 4 - stage 6 | 0.0462 | 0.0096 | 137.2635 | 4.8153 | 0.0002 | *** |
| Stage 5 - stage 6 | -0.0012 | 0.0101 | 133.7009 | -0.1168 | 1.0000 |  |
| Stage 6 - stage 7 | 0.3261 | 0.0381 | 51.6950 | 8.5689 | 0.0001 | *** |
| Stage 7 - stage 8 | 0.1374 | 0.0482 | 29.4633 | 2.8533 | 0.0372 | * |
| Tyramine | estimate | SE | df | t.ratio | p.value | |
| Stage 1 - stage 2 | 0.0031 | 0.0031 | 135.1796 | 1.0269 | 0.9968 |  |
| Stage 1 - stage 3 | 0.0025 | 0.0045 | 17.6703 | 0.5606 | 1.0000 |  |
| Stage 1 - stage 4 | -0.0064 | 0.0046 | 19.8186 | -1.3759 | 0.9555 |  |
| Stage 1 - stage 5 | 0.0129 | 0.0049 | 24.7723 | 2.6203 | 0.3205 |  |
| Stage 1 - stage 6 | 0.0070 | 0.0056 | 38.6179 | 1.2596 | 0.9795 |  |
| Stage 2 - stage 3 | 0.0006 | 0.0045 | 18.2171 | 0.1342 | 1.0000 |  |
| Stage 2 - stage 4 | -0.0032 | 0.0047 | 20.3970 | -0.6945 | 0.9998 |  |
| Stage 2 - stage 5 | -0.0098 | 0.0050 | 25.4058 | -1.9715 | 0.7068 |  |
| Stage 2 - stage 6 | -0.0039 | 0.0056 | 39.3510 | -0.6938 | 0.9999 |  |
| Stage 3 - stage 4 | -0.0039 | 0.0026 | 141.6121 | -1.5086 | 0.9360 |  |
| Stage 3 - stage 5 | 0.0104 | 0.0031 | 143.0217 | 3.3357 | 0.0482 | * |
| Stage 3 - stage 6 | 0.0045 | 0.0041 | 141.1092 | 1.1098 | 0.9938 |  |
| Stage 4 - stage 5 | 0.0065 | 0.0031 | 136.4016 | 2.0816 | 0.6366 |  |
| Stage 4 - stage 6 | 0.0006 | 0.0041 | 137.7285 | 0.1564 | 1.0000 |  |
| Stage 5 - stage 6 | -0.0059 | 0.0044 | 135.9291 | -1.3477 | 0.9710 |  |
| Stage 6 - stage 7 | 0.0487 | 0.0089 | 67.2207 | 5.4612 | 0.0001 | *** |
| Stage 7 - stage 8 | 0.0186 | 0.0105 | 29.9190 | 1.7722 | 0.3062 |  |
| Serotonin | estimate | SE | df | t.ratio | p.value | |
| Stage 1 - stage 2 | 0.0136 | 0.0296 | 85.4927 | 0.4575 | 1.0000 |  |
| Stage 1 - stage 3 | -0.0608 | 0.0477 | 29.9788 | -1.2728 | 0.9766 |  |
| Stage 1 - stage 4 | 0.0420 | 0.0437 | 21.5842 | 0.9606 | 0.9972 |  |
| Stage 1 - stage 5 | 0.0424 | 0.0539 | 43.4003 | 0.7865 | 0.9997 |  |
| Stage 1 - stage 6 | 0.5913 | 0.0538 | 42.8065 | 10.9821 | 0.0001 | *** |
| Stage 2 - stage 3 | -0.0284 | 0.0451 | 24.1479 | -0.6307 | 0.9999 |  |
| Stage 2 - stage 4 | -0.0472 | 0.0490 | 32.6983 | -0.9634 | 0.9976 |  |
| Stage 2 - stage 5 | -0.0288 | 0.0550 | 45.9839 | -0.5239 | 1.0000 |  |
| Stage 2 - stage 6 | -0.5777 | 0.0550 | 45.3799 | -10.5118 | 0.0001 | *** |
| Stage 3 - stage 4 | -0.0188 | 0.0310 | 90.6319 | -0.6049 | 1.0000 |  |
| Stage 3 - stage 5 | 0.0004 | 0.0396 | 90.0450 | 0.0094 | 1.0000 |  |
| Stage 3 - stage 6 | 0.5493 | 0.0400 | 91.9563 | 13.7432 | 0.0001 | *** |
| Stage 4 - stage 5 | -0.0184 | 0.0424 | 86.8150 | -0.4334 | 1.0000 |  |
| Stage 4 - stage 6 | 0.5305 | 0.0458 | 90.9047 | 11.5921 | 0.0001 | *** |
| Stage 5 - stage 6 | 0.5489 | 0.0521 | 89.9877 | 10.5370 | 0.0001 | *** |
| Stage 6 - stage 7 | -0.2756 | 0.1189 | 90.7031 | -2.3176 | 0.6121 |  |
| Stage 7 - stage 8 | 0.3506 | 0.1697 | 23.9792 | 2.0657 | 0.1930 |  |
| Tryptophan | estimate | SE | df | t.ratio | p.value | |
| Stage 1 - stage 2 | 0.0390 | 0.3738 | 153.2773 | 0.1044 | 1.0000 |  |
| Stage 1 - stage 3 | -3.2024 | 0.8449 | 15.3034 | -3.7902 | 0.0517 | * |
| Stage 1 - stage 4 | 1.3605 | 0.8287 | 14.1617 | 1.6417 | 0.8673 |  |
| Stage 1 - stage 5 | 2.2181 | 0.8738 | 17.4311 | 2.5385 | 0.3782 |  |
| Stage 1 - stage 6 | 2.3965 | 0.9410 | 23.1600 | 2.5467 | 0.3611 |  |
| Stage 2 - stage 3 | -1.3215 | 0.8310 | 14.3211 | -1.5903 | 0.8875 |  |
| Stage 2 - stage 4 | -3.1634 | 0.8471 | 15.4687 | -3.7343 | 0.0566 | * |
| Stage 2 - stage 5 | -2.1791 | 0.8759 | 17.6061 | -2.4878 | 0.4042 |  |
| Stage 2 - stage 6 | -2.3575 | 0.9430 | 23.3568 | -2.5000 | 0.3859 |  |
| Stage 3 - stage 4 | -1.8419 | 0.3311 | 155.8332 | -5.5630 | 0.0001 | *** |
| Stage 3 - stage 5 | 0.8576 | 0.4058 | 156.6401 | 2.1133 | 0.6141 |  |
| Stage 3 - stage 6 | 1.0360 | 0.5350 | 155.7889 | 1.9365 | 0.7345 |  |
| Stage 4 - stage 5 | -0.9843 | 0.4194 | 153.9073 | -2.3469 | 0.4485 |  |
| Stage 4 - stage 6 | -0.8059 | 0.5530 | 154.4181 | -1.4572 | 0.9495 |  |
| Stage 5 - stage 6 | 0.1784 | 0.5821 | 153.5676 | 0.3064 | 1.0000 |  |
| Stage 6 - stage 7 | -1.8985 | 1.0141 | 59.7509 | -1.8721 | 0.8779 |  |
| Stage 7 - stage 8 | 1.5080 | 0.6623 | 22.0000 | 2.2769 | 0.1342 |  |
| Tryptamine | estimate | SE | df | t.ratio | p.value | |
| Stage 1 - stage 2 | 0.0109 | 0.0125 | 117.6303 | 0.8710 | 0.9993 |  |
| Stage 1 - stage 3 | 0.0146 | 0.0149 | 29.3669 | 0.9817 | 0.9970 |  |
| Stage 1 - stage 4 | -0.0213 | 0.0147 | 27.3582 | -1.4485 | 0.9423 |  |
| Stage 1 - stage 5 | -0.0201 | 0.0163 | 39.2910 | -1.2299 | 0.9830 |  |
| Stage 1 - stage 6 | -0.0159 | 0.0183 | 55.0954 | -0.8643 | 0.9992 |  |
| Stage 2 - stage 3 | 0.0322 | 0.0140 | 22.8939 | 2.3073 | 0.4987 |  |
| Stage 2 - stage 4 | 0.0255 | 0.0141 | 24.7450 | 1.8049 | 0.8008 |  |
| Stage 2 - stage 5 | 0.0310 | 0.0157 | 34.5698 | 1.9788 | 0.7036 |  |
| Stage 2 - stage 6 | 0.0268 | 0.0177 | 50.6491 | 1.5077 | 0.9315 |  |
| Stage 3 - stage 4 | -0.0067 | 0.0092 | 121.9310 | -0.7330 | 0.9999 |  |
| Stage 3 - stage 5 | 0.0012 | 0.0117 | 123.9014 | 0.1040 | 1.0000 |  |
| Stage 3 - stage 6 | 0.0055 | 0.0142 | 122.1571 | 0.3841 | 1.0000 |  |
| Stage 4 - stage 5 | -0.0055 | 0.0114 | 117.7812 | -0.4815 | 1.0000 |  |
| Stage 4 - stage 6 | -0.0012 | 0.0143 | 119.0582 | -0.0871 | 1.0000 |  |
| Stage 5 - stage 6 | 0.0042 | 0.0155 | 116.6436 | 0.2735 | 1.0000 |  |
| Stage 6 - stage 7 | 0.0124 | 0.0193 | 85.4670 | 0.6429 | 1.0000 |  |
| Stage 7 - stage 8 | 0.0061 | 0.0175 | 32.2935 | 0.3478 | 0.9853 |  |

**Table s6: Hawthorn race developmental stage comparisons for each neurotransmitter.**

| Histidine | estimate | SE | df | t.ratio | p.value | |
| --- | --- | --- | --- | --- | --- | --- |
| Stage 1 - stage 2 | 9.2760 | 3.3472 | 150.2634 | 2.7713 | 0.2033 |  |
| Stage 1 - stage 3 | -8.1420 | 3.6374 | 35.6718 | -2.2384 | 0.5348 |  |
| Stage 1 - stage 4 | -18.5680 | 4.2215 | 57.4697 | -4.3984 | 0.0026 | ** |
| Stage 1 - stage 5 | 1.4017 | 4.9156 | 79.2112 | 0.2852 | 1.0000 |  |
| Stage 1 - stage 6 | -4.5546 | 4.6661 | 62.2434 | -0.9761 | 0.9977 |  |
| Stage 2 - stage 3 | 17.4180 | 3.4578 | 29.7771 | 5.0373 | 0.0011 | ** |
| Stage 2 - stage 4 | -9.2919 | 4.0678 | 51.2306 | -2.2843 | 0.5002 |  |
| Stage 2 - stage 5 | 7.8743 | 4.7842 | 73.8749 | 1.6459 | 0.8858 |  |
| Stage 2 - stage 6 | 13.8306 | 4.5274 | 56.8971 | 3.0548 | 0.1196 |  |
| Stage 3 - stage 4 | -26.7099 | 3.2232 | 160.6241 | -8.2867 | 0.0001 | *** |
| Stage 3 - stage 5 | 9.5437 | 4.1790 | 155.8926 | 2.2837 | 0.4926 |  |
| Stage 3 - stage 6 | 3.5874 | 3.9071 | 134.5693 | 0.9182 | 0.9988 |  |
| Stage 4 - stage 5 | -17.1662 | 4.4386 | 154.5738 | -3.8675 | 0.0086 | ** |
| Stage 4 - stage 6 | -23.1225 | 4.1837 | 160.9020 | -5.5268 | 0.0001 | *** |
| Stage 5 - stage 6 | -5.9563 | 4.7049 | 152.9116 | -1.2660 | 0.9821 |  |
| Stage 6 - stage 7 | 7.4986 | 4.6068 | 75.3566 | 1.6277 | 0.9579 |  |
| Stage 7 - stage 8 | -9.4681 | 1.9680 | 35.4192 | -4.8110 | 0.0002 | *** |
| Serine | estimate | SE | df | t.ratio | p.value | |
| Stage 1 - stage 2 | -1.4183 | 1.8679 | 141.2302 | -0.7593 | 0.9998 |  |
| Stage 1 - stage 3 | -5.3565 | 2.1221 | 23.4911 | -2.5242 | 0.3726 |  |
| Stage 1 - stage 4 | -10.0340 | 2.4487 | 39.8930 | -4.0976 | 0.0094 | ** |
| Stage 1 - stage 5 | 13.0445 | 2.7836 | 53.8388 | 4.6863 | 0.0011 | ** |
| Stage 1 - stage 6 | 0.2133 | 2.7264 | 46.6659 | 0.0782 | 1.0000 |  |
| Stage 2 - stage 3 | 3.9382 | 2.0498 | 20.8278 | 1.9212 | 0.7351 |  |
| Stage 2 - stage 4 | -11.4523 | 2.3864 | 36.8641 | -4.7990 | 0.0014 | ** |
| Stage 2 - stage 5 | -14.4628 | 2.7289 | 51.0300 | -5.2999 | 0.0001 | *** |
| Stage 2 - stage 6 | -1.6316 | 2.6706 | 43.9618 | -0.6110 | 1.0000 |  |
| Stage 3 - stage 4 | -15.3905 | 1.8540 | 152.4109 | -8.3011 | 0.0001 | *** |
| Stage 3 - stage 5 | 18.4009 | 2.3417 | 144.0862 | 7.8580 | 0.0001 | *** |
| Stage 3 - stage 6 | 5.5698 | 2.2836 | 129.6390 | 2.4391 | 0.3880 |  |
| Stage 4 - stage 5 | 3.0104 | 2.4795 | 144.9630 | 1.2141 | 0.9871 |  |
| Stage 4 - stage 6 | -9.8207 | 2.4253 | 150.4528 | -4.0493 | 0.0045 | ** |
| Stage 5 - stage 6 | -12.8311 | 2.6298 | 139.1037 | -4.8790 | 0.0002 | *** |
| Stage 6 - stage 7 | 9.5102 | 3.0334 | 54.4169 | 3.1352 | 0.1522 |  |
| Stage 7 - stage 8 | -5.6389 | 2.1560 | 34.5884 | -2.6154 | 0.0603 |  |
| Histamine | estimate | SE | df | t.ratio | p.value | |
| Stage 1 - stage 2 | 0.0030 | 0.0045 | 121.9995 | 0.6518 | 1.0000 |  |
| Stage 1 - stage 3 | 0.0059 | 0.0068 | 16.9106 | 0.8617 | 0.9987 |  |
| Stage 1 - stage 4 | -0.0244 | 0.0071 | 20.3297 | -3.4219 | 0.0818 |  |
| Stage 1 - stage 5 | 0.0230 | 0.0077 | 26.6557 | 2.9844 | 0.1684 |  |
| Stage 1 - stage 6 | 0.0986 | 0.0077 | 25.6194 | 12.8708 | 0.0001 | *** |
| Stage 2 - stage 3 | -0.0029 | 0.0068 | 16.9128 | -0.4270 | 1.0000 |  |
| Stage 2 - stage 4 | -0.0215 | 0.0071 | 20.3318 | -3.0085 | 0.1759 |  |
| Stage 2 - stage 5 | -0.0201 | 0.0077 | 26.6575 | -2.6018 | 0.3267 |  |
| Stage 2 - stage 6 | -0.0956 | 0.0077 | 25.6213 | -12.4848 | 0.0001 | *** |
| Stage 3 - stage 4 | -0.0186 | 0.0041 | 128.2996 | -4.5462 | 0.0007 | *** |
| Stage 3 - stage 5 | 0.0172 | 0.0052 | 130.5207 | 3.3064 | 0.0532 |  |
| Stage 3 - stage 6 | 0.0927 | 0.0051 | 131.9887 | 18.0226 | 0.0001 | *** |
| Stage 4 - stage 5 | -0.0014 | 0.0051 | 123.1360 | -0.2740 | 1.0000 |  |
| Stage 4 - stage 6 | 0.0741 | 0.0051 | 125.2129 | 14.5311 | 0.0001 | *** |
| Stage 5 - stage 6 | 0.0755 | 0.0056 | 122.7340 | 13.5327 | 0.0001 | *** |
| Stage 6 - stage 7 | 0.4911 | 0.0325 | 42.5499 | 15.1207 | 0.0001 | *** |
| Stage 7 - stage 8 | 0.2389 | 0.0455 | 34.8184 | 5.2453 | 0.0001 | *** |
| Aspartate | estimate | SE | df | t.ratio | p.value | |
| Stage 1 - stage 2 | -1.4488 | 3.7653 | 148.7253 | -0.3848 | 1.0000 |  |
| Stage 1 - stage 3 | 2.6594 | 7.0984 | 13.3624 | 0.3747 | 1.0000 |  |
| Stage 1 - stage 4 | -51.1903 | 7.4971 | 16.5096 | -6.8280 | 0.0002 | *** |
| Stage 1 - stage 5 | 30.0268 | 7.9971 | 20.9307 | 3.7547 | 0.0409 | * |
| Stage 1 - stage 6 | 25.4946 | 7.9695 | 20.4891 | 3.1990 | 0.1244 |  |
| Stage 2 - stage 3 | -4.1083 | 7.0651 | 13.1084 | -0.5815 | 1.0000 |  |
| Stage 2 - stage 4 | -52.6391 | 7.4656 | 16.2307 | -7.0509 | 0.0001 | *** |
| Stage 2 - stage 5 | -31.4756 | 7.9675 | 20.6243 | -3.9505 | 0.0275 | * |
| Stage 2 - stage 6 | -26.9434 | 7.9398 | 20.1867 | -3.3934 | 0.0868 |  |
| Stage 3 - stage 4 | -48.5308 | 3.7572 | 158.0238 | -12.9167 | 0.0001 | *** |
| Stage 3 - stage 5 | 27.3673 | 4.8379 | 159.3046 | 5.6568 | 0.0001 | *** |
| Stage 3 - stage 6 | 22.8352 | 4.8331 | 160.4201 | 4.7247 | 0.0003 | *** |
| Stage 4 - stage 5 | -21.1635 | 4.7285 | 149.9532 | -4.4757 | 0.0009 | *** |
| Stage 4 - stage 6 | -25.6957 | 4.7262 | 152.1083 | -5.4369 | 0.0001 | *** |
| Stage 5 - stage 6 | -4.5322 | 5.1331 | 149.2986 | -0.8829 | 0.9992 |  |
| Stage 6 - stage 7 | 138.0729 | 14.7200 | 23.1542 | 9.3800 | 0.0001 | *** |
| Stage 7 - stage 8 | 1.9889 | 11.9196 | 34.6003 | 0.1669 | 0.9983 |  |
| Glutamate | estimate | SE | df | t.ratio | p.value | |
| Stage 1 - stage 2 | 10.2681 | 7.1871 | 150.0784 | 1.4287 | 0.9560 |  |
| Stage 1 - stage 3 | -9.7917 | 9.4441 | 20.5562 | -1.0368 | 0.9945 |  |
| Stage 1 - stage 4 | -54.0539 | 10.4326 | 29.4495 | -5.1813 | 0.0008 | *** |
| Stage 1 - stage 5 | -4.7870 | 11.5635 | 40.6643 | -0.4140 | 1.0000 |  |
| Stage 1 - stage 6 | -5.9932 | 11.4700 | 38.2213 | -0.5225 | 1.0000 |  |
| Stage 2 - stage 3 | 20.0598 | 9.4475 | 20.5660 | 2.1233 | 0.6138 |  |
| Stage 2 - stage 4 | -43.7859 | 10.4356 | 29.4562 | -4.1958 | 0.0102 | * |
| Stage 2 - stage 5 | 15.0551 | 11.5662 | 40.6661 | 1.3016 | 0.9742 |  |
| Stage 2 - stage 6 | 16.2612 | 11.4728 | 38.2242 | 1.4174 | 0.9527 |  |
| Stage 3 - stage 4 | -63.8456 | 6.7773 | 160.6880 | -9.4205 | 0.0001 | *** |
| Stage 3 - stage 5 | 5.0047 | 8.6549 | 160.9555 | 0.5782 | 1.0000 |  |
| Stage 3 - stage 6 | 3.7985 | 8.5995 | 158.9816 | 0.4417 | 1.0000 |  |
| Stage 4 - stage 5 | -58.8410 | 8.8143 | 151.6863 | -6.6757 | 0.0001 | *** |
| Stage 4 - stage 6 | -60.0471 | 8.7713 | 155.7959 | -6.8459 | 0.0001 | *** |
| Stage 5 - stage 6 | -1.2061 | 9.5939 | 151.2755 | -0.1257 | 1.0000 |  |
| Stage 6 - stage 7 | 130.7025 | 13.1765 | 45.8484 | 9.9194 | 0.0001 | *** |
| Stage 7 - stage 8 | 0.5994 | 10.5082 | 32.8034 | 0.0570 | 0.9999 |  |
| GABA | estimate | SE | df | t.ratio | p.value | |
| Stage 1 - stage 2 | 0.4085 | 0.6457 | 146.4839 | 0.6327 | 1.0000 |  |
| Stage 1 - stage 3 | -0.1254 | 0.6311 | 42.5976 | -0.1987 | 1.0000 |  |
| Stage 1 - stage 4 | 2.1235 | 0.7393 | 66.2943 | 2.8722 | 0.1752 |  |
| Stage 1 - stage 5 | 4.5432 | 0.8546 | 82.2229 | 5.3160 | 0.0001 | *** |
| Stage 1 - stage 6 | 14.0671 | 0.8636 | 75.1907 | 16.2892 | 0.0001 | *** |
| Stage 2 - stage 3 | 0.5339 | 0.6300 | 42.8572 | 0.8475 | 0.9993 |  |
| Stage 2 - stage 4 | 2.5320 | 0.7384 | 66.7022 | 3.4292 | 0.0446 | * |
| Stage 2 - stage 5 | -4.1347 | 0.8538 | 82.6405 | -4.8428 | 0.0004 | *** |
| Stage 2 - stage 6 | -13.6586 | 0.8628 | 75.5517 | -15.8314 | 0.0001 | *** |
| Stage 3 - stage 4 | 1.9981 | 0.5893 | 152.0020 | 3.3909 | 0.0405 | * |
| Stage 3 - stage 5 | 4.6685 | 0.7418 | 138.7799 | 6.2935 | 0.0001 | *** |
| Stage 3 - stage 6 | 14.1924 | 0.7563 | 120.7116 | 18.7656 | 0.0001 | *** |
| Stage 4 - stage 5 | 6.6667 | 0.7915 | 152.1419 | 8.4226 | 0.0001 | *** |
| Stage 4 - stage 6 | 16.1906 | 0.8059 | 157.6172 | 20.0903 | 0.0001 | *** |
| Stage 5 - stage 6 | 9.5239 | 0.8866 | 150.5125 | 10.7419 | 0.0001 | *** |
| Stage 6 - stage 7 | 44.5772 | 2.8134 | 31.8033 | 15.8444 | 0.0001 | *** |
| Stage 7 - stage 8 | 0.1193 | 3.1720 | 32.6319 | 0.0376 | 1.0000 |  |
| Dopa | estimate | SE | df | t.ratio | p.value | |
| Stage 1 - stage 2 | -0.0152 | 0.0260 | 149.2417 | -0.5864 | 1.0000 |  |
| Stage 1 - stage 3 | -0.0574 | 0.0259 | 50.5113 | -2.2141 | 0.5476 |  |
| Stage 1 - stage 4 | -0.1969 | 0.0317 | 84.6373 | -6.2059 | 0.0001 | *** |
| Stage 1 - stage 5 | -0.0065 | 0.0353 | 90.9806 | -0.1840 | 1.0000 |  |
| Stage 1 - stage 6 | -0.0008 | 0.0356 | 85.1021 | -0.0212 | 1.0000 |  |
| Stage 2 - stage 3 | 0.0421 | 0.0249 | 45.0092 | 1.6911 | 0.8628 |  |
| Stage 2 - stage 4 | -0.2121 | 0.0309 | 80.2057 | -6.8597 | 0.0001 | *** |
| Stage 2 - stage 5 | -0.0087 | 0.0346 | 87.4519 | -0.2521 | 1.0000 |  |
| Stage 2 - stage 6 | -0.0145 | 0.0349 | 81.6114 | -0.4143 | 1.0000 |  |
| Stage 3 - stage 4 | -0.2542 | 0.0258 | 154.8789 | -9.8450 | 0.0001 | *** |
| Stage 3 - stage 5 | 0.0509 | 0.0307 | 137.2628 | 1.6545 | 0.8851 |  |
| Stage 3 - stage 6 | 0.0566 | 0.0313 | 123.8106 | 1.8118 | 0.8085 |  |
| Stage 4 - stage 5 | -0.2034 | 0.0341 | 153.5463 | -5.9646 | 0.0001 | *** |
| Stage 4 - stage 6 | -0.1976 | 0.0345 | 157.0370 | -5.7267 | 0.0001 | *** |
| Stage 5 - stage 6 | 0.0057 | 0.0368 | 151.8266 | 0.1559 | 1.0000 |  |
| Stage 6 - stage 7 | -0.0487 | 0.0338 | 105.8274 | -1.4395 | 0.9865 |  |
| Stage 7 - stage 8 | -0.0060 | 0.0017 | 33.1420 | -3.5825 | 0.0057 | ** |
| Octopamine | estimate | SE | df | t.ratio | p.value | |
| Stage 1 - stage 2 | 0.0006 | 0.0447 | 78.2365 | 0.0131 | 1.0000 |  |
| Stage 1 - stage 3 | 0.0139 | 0.0604 | 16.4742 | 0.2296 | 1.0000 |  |
| Stage 1 - stage 4 | -0.0067 | 0.0600 | 16.1625 | -0.1125 | 1.0000 |  |
| Stage 1 - stage 5 | 0.0112 | 0.0653 | 21.5997 | 0.1722 | 1.0000 |  |
| Stage 1 - stage 6 | 0.0916 | 0.0636 | 19.3057 | 1.4405 | 0.9400 |  |
| Stage 2 - stage 3 | -0.0133 | 0.0568 | 13.2860 | -0.2337 | 1.0000 |  |
| Stage 2 - stage 4 | -0.0107 | 0.0620 | 18.1907 | -0.1718 | 1.0000 |  |
| Stage 2 - stage 5 | -0.0910 | 0.0602 | 16.0158 | -1.5113 | 0.9168 |  |
| Stage 2 - stage 6 | -0.0062 | 0.0564 | 12.9841 | -0.1092 | 1.0000 |  |
| Stage 3 - stage 4 | 0.0071 | 0.0370 | 81.0528 | 0.1925 | 1.0000 |  |
| Stage 3 - stage 5 | -0.0026 | 0.0463 | 84.3525 | -0.0565 | 1.0000 |  |
| Stage 3 - stage 6 | 0.0777 | 0.0446 | 86.9104 | 1.7414 | 0.8437 |  |
| Stage 4 - stage 5 | 0.0045 | 0.0434 | 79.3577 | 0.1037 | 1.0000 |  |
| Stage 4 - stage 6 | 0.0848 | 0.0420 | 82.5509 | 2.0209 | 0.6781 |  |
| Stage 5 - stage 6 | 0.0803 | 0.0478 | 81.8784 | 1.6801 | 0.8719 |  |
| Stage 6 - stage 7 | 0.2462 | 0.0482 | 37.7643 | 5.1098 | 0.0009 | *** |
| Stage 7 - stage 8 | 0.1023 | 0.0294 | 34.0988 | 3.4782 | 0.0073 | ** |
| Tyrosine | estimate | SE | df | t.ratio | p.value | |
| Stage 1 - stage 2 | -0.1615 | 3.2025 | 143.9551 | -0.0504 | 1.0000 |  |
| Stage 1 - stage 3 | 0.8927 | 3.6657 | 20.3734 | 0.2435 | 1.0000 |  |
| Stage 1 - stage 4 | -59.8429 | 4.2644 | 33.6500 | -14.0332 | 0.0001 | *** |
| Stage 1 - stage 5 | 41.4830 | 4.7469 | 44.5304 | 8.7390 | 0.0001 | *** |
| Stage 1 - stage 6 | 32.0165 | 4.6897 | 39.2125 | 6.8270 | 0.0001 | *** |
| Stage 2 - stage 3 | -1.0542 | 3.6144 | 19.2515 | -0.2917 | 1.0000 |  |
| Stage 2 - stage 4 | -60.0044 | 4.2204 | 32.3473 | -14.2178 | 0.0001 | *** |
| Stage 2 - stage 5 | -41.6445 | 4.7074 | 43.2119 | -8.8467 | 0.0001 | *** |
| Stage 2 - stage 6 | -32.1780 | 4.6497 | 37.9893 | -6.9204 | 0.0001 | *** |
| Stage 3 - stage 4 | -58.9502 | 3.2072 | 149.0144 | -18.3804 | 0.0001 | *** |
| Stage 3 - stage 5 | 40.5903 | 3.9062 | 143.0938 | 10.3914 | 0.0001 | *** |
| Stage 3 - stage 6 | 31.1238 | 3.8641 | 123.5099 | 8.0546 | 0.0001 | *** |
| Stage 4 - stage 5 | -18.3599 | 4.0528 | 146.7099 | -4.5302 | 0.0007 | *** |
| Stage 4 - stage 6 | -27.8264 | 4.0274 | 155.4292 | -6.9092 | 0.0001 | *** |
| Stage 5 - stage 6 | -9.4665 | 4.3606 | 147.8187 | -2.1709 | 0.5731 |  |
| Stage 6 - stage 7 | -25.1479 | 4.4380 | 57.7244 | -5.6665 | 0.0001 | *** |
| Stage 7 - stage 8 | -5.0456 | 1.0150 | 34.7425 | -4.9710 | 0.0001 | *** |
| Dopamine | estimate | SE | df | t.ratio | p.value | |
| Stage 1 - stage 2 | 0.0179 | 0.0064 | 130.3156 | 2.7912 | 0.1962 |  |
| Stage 1 - stage 3 | -0.0119 | 0.0070 | 43.8055 | -1.6919 | 0.8622 |  |
| Stage 1 - stage 4 | 0.0030 | 0.0079 | 59.5628 | 0.3819 | 1.0000 |  |
| Stage 1 - stage 5 | 0.0026 | 0.0091 | 76.0666 | 0.2900 | 1.0000 |  |
| Stage 1 - stage 6 | 0.0243 | 0.0091 | 73.0937 | 2.6703 | 0.2620 |  |
| Stage 2 - stage 3 | 0.0297 | 0.0070 | 43.9311 | 4.2419 | 0.0056 | ** |
| Stage 2 - stage 4 | 0.0209 | 0.0078 | 59.7562 | 2.6598 | 0.2716 |  |
| Stage 2 - stage 5 | 0.0152 | 0.0091 | 76.2884 | 1.6825 | 0.8705 |  |
| Stage 2 - stage 6 | -0.0064 | 0.0091 | 73.3004 | -0.7074 | 0.9999 |  |
| Stage 3 - stage 4 | -0.0089 | 0.0066 | 138.7670 | -1.3402 | 0.9722 |  |
| Stage 3 - stage 5 | 0.0145 | 0.0082 | 136.8448 | 1.7644 | 0.8341 |  |
| Stage 3 - stage 6 | 0.0362 | 0.0083 | 133.0699 | 4.3522 | 0.0015 | ** |
| Stage 4 - stage 5 | 0.0056 | 0.0085 | 132.1090 | 0.6639 | 0.9999 |  |
| Stage 4 - stage 6 | 0.0273 | 0.0086 | 136.8735 | 3.1653 | 0.0782 |  |
| Stage 5 - stage 6 | 0.0217 | 0.0095 | 132.5954 | 2.2841 | 0.4930 |  |
| Stage 6 - stage 7 | 0.4249 | 0.0366 | 43.3556 | 11.6190 | 0.0001 | *** |
| Stage 7 - stage 8 | -0.0389 | 0.0497 | 29.5559 | -0.7835 | 0.8613 |  |
| Tyramine | estimate | SE | df | t.ratio | p.value | |
| Stage 1 - stage 2 | 0.0004 | 0.0030 | 135.9730 | 0.1404 | 1.0000 |  |
| Stage 1 - stage 3 | 0.0034 | 0.0045 | 17.5711 | 0.7463 | 0.9997 |  |
| Stage 1 - stage 4 | -0.0255 | 0.0048 | 22.7596 | -5.2865 | 0.0011 | ** |
| Stage 1 - stage 5 | 0.0215 | 0.0052 | 28.9461 | 4.1524 | 0.0116 | * |
| Stage 1 - stage 6 | 0.0242 | 0.0053 | 30.0147 | 4.5980 | 0.0035 | ** |
| Stage 2 - stage 3 | -0.0029 | 0.0044 | 15.5702 | -0.6743 | 0.9999 |  |
| Stage 2 - stage 4 | -0.0251 | 0.0047 | 20.5446 | -5.3425 | 0.0013 | ** |
| Stage 2 - stage 5 | -0.0211 | 0.0051 | 26.5769 | -4.1680 | 0.0124 | * |
| Stage 2 - stage 6 | -0.0238 | 0.0051 | 27.6413 | -4.6222 | 0.0038 | ** |
| Stage 3 - stage 4 | -0.0221 | 0.0027 | 143.6473 | -8.1276 | 0.0001 | *** |
| Stage 3 - stage 5 | 0.0182 | 0.0034 | 146.1464 | 5.2853 | 0.0001 | *** |
| Stage 3 - stage 6 | 0.0208 | 0.0036 | 146.8018 | 5.8266 | 0.0001 | *** |
| Stage 4 - stage 5 | -0.0040 | 0.0035 | 137.0306 | -1.1506 | 0.9916 |  |
| Stage 4 - stage 6 | -0.0013 | 0.0036 | 139.6144 | -0.3674 | 1.0000 |  |
| Stage 5 - stage 6 | 0.0027 | 0.0038 | 136.6062 | 0.6924 | 0.9999 |  |
| Stage 6 - stage 7 | 0.0214 | 0.0080 | 46.5211 | 2.6795 | 0.3735 |  |
| Stage 7 - stage 8 | -0.0045 | 0.0104 | 31.4902 | -0.4360 | 0.9718 |  |
| Serotonin | estimate | SE | df | t.ratio | p.value | |
| Stage 1 - stage 2 | 0.0060 | 0.0273 | 89.8211 | 0.2209 | 1.0000 |  |
| Stage 1 - stage 3 | 0.0562 | 0.0426 | 20.5010 | 1.3176 | 0.9669 |  |
| Stage 1 - stage 4 | -0.0151 | 0.0496 | 31.2669 | -0.3041 | 1.0000 |  |
| Stage 1 - stage 5 | -0.0227 | 0.0508 | 36.3307 | -0.4476 | 1.0000 |  |
| Stage 1 - stage 6 | 0.1396 | 0.0503 | 31.7630 | 2.7759 | 0.2376 |  |
| Stage 2 - stage 3 | -0.0502 | 0.0435 | 21.1722 | -1.1519 | 0.9876 |  |
| Stage 2 - stage 4 | -0.0091 | 0.0504 | 31.6932 | -0.1799 | 1.0000 |  |
| Stage 2 - stage 5 | 0.0288 | 0.0516 | 36.6551 | 0.5577 | 1.0000 |  |
| Stage 2 - stage 6 | -0.1335 | 0.0510 | 32.1716 | -2.6162 | 0.3121 |  |
| Stage 3 - stage 4 | 0.0411 | 0.0383 | 92.3420 | 1.0737 | 0.9951 |  |
| Stage 3 - stage 5 | -0.0789 | 0.0405 | 94.9835 | -1.9471 | 0.7267 |  |
| Stage 3 - stage 6 | 0.0834 | 0.0404 | 91.1039 | 2.0635 | 0.6493 |  |
| Stage 4 - stage 5 | -0.0378 | 0.0419 | 89.7910 | -0.9034 | 0.9989 |  |
| Stage 4 - stage 6 | 0.1245 | 0.0465 | 93.0608 | 2.6755 | 0.2551 |  |
| Stage 5 - stage 6 | 0.1623 | 0.0449 | 94.8561 | 3.6159 | 0.0230 | * |
| Stage 6 - stage 7 | 0.2767 | 0.1106 | 64.5195 | 2.5028 | 0.4824 |  |
| Stage 7 - stage 8 | -0.0828 | 0.1657 | 24.8909 | -0.4998 | 0.9584 |  |
| Tryptophan | estimate | SE | df | t.ratio | p.value | |
| Stage 1 - stage 2 | -0.0291 | 0.3596 | 153.2724 | -0.0808 | 1.0000 |  |
| Stage 1 - stage 3 | 0.7692 | 0.8231 | 13.7971 | 0.9345 | 0.9971 |  |
| Stage 1 - stage 4 | -4.6197 | 0.8570 | 16.1579 | -5.3908 | 0.0024 | ** |
| Stage 1 - stage 5 | 1.9547 | 0.8934 | 18.9130 | 2.1878 | 0.5752 |  |
| Stage 1 - stage 6 | 1.1053 | 0.9029 | 19.6164 | 1.2242 | 0.9800 |  |
| Stage 2 - stage 3 | -0.7982 | 0.8205 | 13.6208 | -0.9729 | 0.9959 |  |
| Stage 2 - stage 4 | -4.6487 | 0.8544 | 15.9689 | -5.4408 | 0.0022 | ** |
| Stage 2 - stage 5 | -1.9837 | 0.8910 | 18.7114 | -2.2264 | 0.5520 |  |
| Stage 2 - stage 6 | -1.1344 | 0.9005 | 19.4122 | -1.2597 | 0.9753 |  |
| Stage 3 - stage 4 | -3.8505 | 0.3574 | 157.1249 | -10.7744 | 0.0001 | *** |
| Stage 3 - stage 5 | 1.1855 | 0.4544 | 158.8217 | 2.6090 | 0.2837 |  |
| Stage 3 - stage 6 | 0.3362 | 0.4766 | 159.9671 | 0.7053 | 0.9999 |  |
| Stage 4 - stage 5 | -2.6650 | 0.4616 | 154.0365 | -5.7735 | 0.0001 | *** |
| Stage 4 - stage 6 | -3.5143 | 0.4825 | 155.3909 | -7.2829 | 0.0001 | *** |
| Stage 5 - stage 6 | -0.8493 | 0.5133 | 153.9906 | -1.6546 | 0.8854 |  |
| Stage 6 - stage 7 | 0.2382 | 0.8662 | 33.6721 | 0.2749 | 1.0000 |  |
| Stage 7 - stage 8 | -1.0817 | 0.6399 | 22.0000 | -1.6905 | 0.3522 |  |
| Tryptamine | estimate | SE | df | t.ratio | p.value | |
| Stage 1 - stage 2 | -0.0121 | 0.0119 | 115.5362 | -1.0167 | 0.9970 |  |
| Stage 1 - stage 3 | -0.0372 | 0.0140 | 23.2468 | -2.6650 | 0.3017 |  |
| Stage 1 - stage 4 | 0.0256 | 0.0148 | 28.2896 | 1.7315 | 0.8391 |  |
| Stage 1 - stage 5 | -0.0396 | 0.0166 | 39.9137 | -2.3893 | 0.4358 |  |
| Stage 1 - stage 6 | -0.0389 | 0.0160 | 34.4596 | -2.4229 | 0.4184 |  |
| Stage 2 - stage 3 | 0.0251 | 0.0137 | 21.5369 | 1.8361 | 0.7826 |  |
| Stage 2 - stage 4 | 0.0135 | 0.0145 | 26.4791 | 0.9301 | 0.9980 |  |
| Stage 2 - stage 5 | 0.0275 | 0.0163 | 38.0354 | 1.6838 | 0.8644 |  |
| Stage 2 - stage 6 | 0.0268 | 0.0158 | 32.6616 | 1.6954 | 0.8576 |  |
| Stage 3 - stage 4 | -0.0116 | 0.0094 | 123.7577 | -1.2323 | 0.9853 |  |
| Stage 3 - stage 5 | -0.0024 | 0.0124 | 123.9744 | -0.1916 | 1.0000 |  |
| Stage 3 - stage 6 | -0.0017 | 0.0118 | 120.8547 | -0.1407 | 1.0000 |  |
| Stage 4 - stage 5 | -0.0140 | 0.0123 | 116.6200 | -1.1395 | 0.9921 |  |
| Stage 4 - stage 6 | -0.0133 | 0.0117 | 120.6759 | -1.1403 | 0.9921 |  |
| Stage 5 - stage 6 | 0.0007 | 0.0132 | 115.8549 | 0.0537 | 1.0000 |  |
| Stage 6 - stage 7 | 0.0738 | 0.0166 | 52.3531 | 4.4421 | 0.0042 | ** |
| Stage 7 - stage 8 | -0.0740 | 0.0167 | 34.5264 | -4.4230 | 0.0005 | *** |

**Table s7: Two-way ANOVA result table showing host race comparisons at stage 4 for selected neurotransmitters.**

| Tyrosine | df | F | P | error |
| --- | --- | --- | --- | --- |
| Hostrace | 1 | 13.231 | 0.005 |  |
| Days | 12 | 4.936 | 0.008 |  |
| Hostrace*Days | 12 | 10.978 | 0.0001 | 10 |
| Tyramine | df | F | P | error |
| Hostrace | 1 | 9.705 | 0.011 |  |
| Days | 12 | 3.153 | 0.039 |  |
| Hostrace*Days | 12 | 4.957 | 0.008 | 10 |
| DOPA | df | F | P | error |
| Hostrace | 1 | 0.649 | 0.447 |  |
| Days | 12 | 0.639 | 0.764 |  |
| Hostrace*Days | 12 | 0.828 | 0.632 | 7 |
| Aspartate | df | F | P | error |
| Hostrace | 1 | 2.082 | 0.177 |  |
| Days | 12 | 4.118 | 0.013 |  |
| Hostrace*Days | 12 | 10.129 | 0.0001 | 11 |
| Glutamate | df | F | P | error |
| Hostrace | 1 | 2.476 | 0.144 |  |
| Days | 12 | 2.918 | 0.043 |  |
| Hostrace*Days | 12 | 6.479 | 0.002 | 11 |
| Tryptophan | df | F | P | error |
| Hostrace | 1 | 1.517 | 0.246 |  |
| Days | 12 | 2.285 | 0.1 |  |
| Hostrace*Days | 12 | 4.359 | 0.013 | 10 |

**Table s8: Two-way ANOVA result table showing host race comparisons at stage 4 for selected neurotransmitters after removing early time point till day20.**

| Tyrosine | df | F | P | error |
| --- | --- | --- | --- | --- |
| Hostrace | 1 | 67.113 | 0.0001 |  |
| Days | 8 | 5.786 | 0.008 |  |
| Hostrace*Days | 8 | 38.149 | 0.0001 | 9 |
| Tyramine | df | F | P | error |
| Hostrace | 1 | 9.261 | 0.012 |  |
| Days | 8 | 2.052 | 0.142 |  |
| Hostrace*Days | 7 | 4.463 | 0.017 | 10 |
| DOPA | df | F | P | error |
| Hostrace | 1 | 0.4 | 0.545 |  |
| Days | 8 | 0.198 | 0.983 |  |
| Hostrace*Days | 7 | 0.846 | 0.581 | 8 |
| Aspartate | df | F | P | error |
| Hostrace | 1 | 5.087 | 0.048 |  |
| Days | 8 | 0.863 | 0.574 |  |
| Hostrace*Days | 8 | 14.25 | 0.0001 | 10 |
| Glutamate | df | F | P | error |
| Hostrace | 1 | 26.94 | 0.001 |  |
| Days | 8 | 4.382 | 0.02 |  |
| Hostrace*Days | 8 | 25.367 | 0.0001 | 9 |
| Tryptophan | df | F | P | error |
| Hostrace | 1 | 1.272 | 0.286 |  |
| Days | 8 | 1.716 | 0.209 |  |
| Hostrace*Days | 7 | 7.193 | 0.003 | 10 |

**Table s9: Instrument method details for liquid chromatography (LC).**

| **Instrument** | Agilent 1290 Infinity UHPLC |
| --- | --- |
| **Column** | Waters Acquity HSS C18, 1.8u, 100mmx2.1mm |
| **Mobile Phase A** | 10mM Ammonium Acetate in Water (0.1%FA) |
| **Mobile Phase B** | Acetonitrile (0.1%FA) |
| **Flow Rate** | 0.2ml/min |
| **Column Oven** | 40°C |
| **Auto-sampler Temp.** | 10°C |
| **Injection Volume** | 10ul |
| **Run Time** | 35mins |
| **Gradient** | 0-3mins:2%B,  3-20mins:2-20%B,  20-25mins:35%B,  25.1mins:80%B,  27mins:80%B,  27.1-35:2%B |

**Table s10: Instrument method details for mass spctrometry (MS).**

| **Instrument** | Thermo Fisher- TSQ Vantage |
| --- | --- |
| **Spray Voltage (+ve)** | 3700V |
| **Vaporizer temp** | 80°C |
| **Sheath gas flow rate** | 20Arb |
| **Aux gas flow rate** | 10Arb |
| **Peak width settings** | 0.70FWHM |
| **Scan time** | 0.05s |
| **Injector settings** | 0-3mins:waste, 3-28.5mins:load, 28.5-35mins:waste |

**Table s11**: Internal standard molecular weights (MW) and expected retention times (RT) for the 14 neurotransmitters and their internal standards used for quantification in mass spectrometry.

| **compound** | **MW** | **RT** |
| --- | --- | --- |
| Histidine-D3 | 329 | 7.53 |
| Histidine | 326.11 | 7.56 |
| serine-D3 | 279.2 | 8.53 |
| Serine | 276.1 | 8.59 |
| Histamine-D4 | 286.37 | 9.33 |
| Histamine | 282.37 | 9.36 |
| Aspartate-D4 | 307.1 | 9.62 |
| Aspartate | 304.1 | 9.68 |
| Glutamate-D5 | 323.26 | 10.48 |
| Glutamate | 318.1 | 10.54 |
| GABA-D6 | 280.32 | 12.96 |
| GABA | 274.28 | 13.06 |
| Dopa-D3 | 371.37 | 15.29 |
| Dopa | 368.35 | 15.33 |
| Octopamine-D3 | 327.17 | 15.44 |
| Octopamine | 324.17 | 15.5 |
| Tyrosine-D6 | 359.4 | 17.25 |
| Tyrosine | 352.2 | 17.35 |
| Dopamine-D4 | 328.37 | 17.85 |
| Dopamine | 324.3 | 17.94 |
| Tyramine-D4 | 312.36 | 20.18 |
| Tyramine | 308.36 | 20.37 |
| Serotonin-D4 | 351.4 | 22.03 |
| Serotonin | 347.38 | 22.11 |
| Tryptophan-D5 | 380.43 | 23.83 |
| Tryptophan | 375.19 | 23.91 |
| Tryptamine-D4 | 335.4 | 26.19 |
| Tryptamine | 331.38 | 26.25 |

**Table s12:** **Standard solution preparation concentrations for all neurotransmitters. ***

|  | Standards | | | Internal standards |
| --- | --- | --- | --- | --- |
| Compound | Pre diapause | Post diapause | Adult fly | ISTD |
| name | on column (ng) | on column (ng) | on column (ng) | on column (ng) |
| Histidine | 20 | 50 | 50 | 1 |
| Serine | 8 | 80 | 80 | 1 |
| Histamine | 3.2 | 3.2 | 3.2 | 1 |
| Aspartate | 25 | 250 | 250 | 1 |
| Glutamate | 25 | 250 | 250 | 1 |
| GABA | 10 | 100 | 100 | 1 |
| Dopa | 0.16 | 0.16 | 0.1 | 1 |
| Octopamine | 1 | 1 | 1 | 1 |
| Tyrosine | 4 | 40 | 40 | 1 |
| Dopamine | 1 | 1 | 1 | 1 |
| Tyramine | 0.32 | 0.32 | 0.64 | 1 |
| Serotonin | 1 | 1 | 1.6 | 1 |
| Tryptophan | 1 | 8 | 8 | 1 |
| Tryptamine | 1 | 1 | 0.16 | 1 |

*The 14 neurotransmitters were weighed individually and standard stock solutions of 1mg/ml in 0.1N HCL were prepared. The final concentration for each chemical for each experiment was calculated by running a preliminary experiment with calibration curves used for honey bee whole brain as in Ramesh et al 2019 and three replicates of apple/ hawthorn brain samples. The internal standard (ISTD) stock solution of 1mg/ml in 0.1NHCL was also prepared.

**Movie 1: Confocal Z-projection stacks (1 µm slices, 100 µm total depth) showing nc82 staining of the pre-diapause larval brain (stage 1, Figure 1).**

**Movie 2: Confocal Z-projection stacks (1 µm slices, 100 µm total depth) showing nc82 staining of the early diapause larval brain (stage 2, Figure 1).**

**Movie 3: Confocal Z-projection stacks (1 µm slices, 100 µm total depth) showing nc82 staining of the late diapause larval brain (stage 3, Figure 1).**

**Movie 4: Confocal Z-projection stacks (1 µm slices, 100 µm total depth) showing nc82 staining of the diapause termination stage and the early adult brain (stage 4, Figure 1).**

**Movie 5: Confocal Z-projection stacks (1 µm slices, 100 µm total depth) showing nc82 staining of developing adult brain (stage 5, Figure 1).**

**Movie 6: Confocal Z-projection stacks (1 µm slices, 100 µm total depth) showing nc82 staining of the late developing adult brain (stage 6, Figure 1).**

**Movie 7: Confocal Z-projection stacks (1 µm slices, 100 µm total depth) showing nc82 staining of the eclosed adult fly brain (stage 7, Figure 1).**
